# Glycosaminoglycan presentation-dependent regulation of glial responses by Orion during *Drosophila* neuronal pruning

**DOI:** 10.64898/2026.09.29.755054

**Authors:** Charline Gal, Clarisse Perron, Sylvain D. Vallet, Evelyne Gout, Margot Weber, Oulidi Fatima Ezzahra, Hui Ji, Chun Han, Hugues Lortat-Jacob, Jean-Maurice Dura, Ana Boulanger

## Abstract

Neuronal pruning refines neural circuits by eliminating unnecessary connections to ensure proper nervous system function. In *Drosophila* mushroom body γ-neuron remodeling, the chemokine-like protein Orion is required for astrocyte infiltration and phagocytosis. This study investigated whether glial responses depend on how Orion is presented. We found that Orion interacts with glycosaminoglycans (GAG) associated to proteoglycans, which are essential for proper axon pruning. Tethering Orion to the γ-neuron plasma membrane, mimicking glycosaminoglycan binding, was sufficient to induce astrocyte infiltration into axon bundles. However, membrane-bound Orion alone could not efficiently support phagocytosis, which additionally requires a soluble and mobile Orion pool. We also identified exosomes carrying GAG-bound Orion as potential mediators of neuronal remodeling. These findings provide the first evidence that glial infiltration can be uncoupled from phagocytosis, demonstrating that the mode of Orion presentation regulates distinct glial mechanisms during neuronal remodeling.

**Significance Statement:** Neuron-glia interactions are critical for developmental neuronal remodeling, an evolutionarily conserved process that shapes neural circuits. During remodeling, glial cells infiltrate degenerating axons and engulf neuronal debris. However, the molecular and cellular mechanisms required for both glial infiltration and phagocytosis remain unknown. This study demonstrates that distinct modes of ligand presentation of the *Drosophila* chemokine-like protein Orion, associated or not with plasma membrane glycosaminoglycans, uncouple infiltration from phagocytosis during remodeling. Because Orion signals through the receptor Draper (the homolog of vertebrate MEGF10) this discovery extends beyond *Drosophila*. Furthermore, it provides a new evolutionary framework for how ligand presentation differentially regulates glial responses. These findings offer crucial insights into neuron-glia communication across species during nervous system development, remodeling, and neurodegeneration.

## Introduction

During development throughout the animal kingdom, neurons initially establish exuberant and often inappropriate axonal and dendritic connections, which are subsequently remodeled during a critical later developmental window. Notably, similar molecular and cellular mechanisms also operate in neurodevelopmental disorders and following nervous system injury ^1, 2, 3, 4, 5, 6^. Neural remodeling generates large amounts of neuronal debris that must be efficiently cleared. Glial cells are the primary phagocytes responsible for debris removal during remodeling, and recent studies have identified key neuron-glia signaling pathways regulating this process ^7^. The elimination of neuronal debris by glial cells can be divided into three main steps. First, glial cells infiltrate axon bundles to access the neuronal material targeted for degradation. Second, they recognize and engulf the debris. Third, the engulfed material is phagocytosed, forming phagosomes that mature to complete phagocytosis.

In *Drosophila*, mushroom bodies (MBs), involved in olfactory learning and memory, undergo extensive remodeling during metamorphosis ^8, 9, 10, 11^. Fragmentation of γ axons is driven by an intrinsic genetic program within MB γ neurons, which critically depends on tightly controlled transcriptional regulation of the ecdysone receptor ^8, 12^. Additionally, astrocytic-like glia (designated here as astrocytes) surrounding the MB play an essential role in remodeling; blocking their infiltration into the MBs prevents this process ^13, 14, 15, 16^. Thus, the infiltration of astrocytic processes into the MB γ axon bundle appears to be a crucial initial step in the glial-mediated elimination of these axons.

Little is known about neuron-secreted ligands that activate glial cells and allow the identification of neuronal targets. In vertebrates, chemokine signaling is one such pathway, crucial for guiding glial migration during development and after injury ^16, 17^. We identified a chemokine-like ligand secreted by MB γ neurons, named Orion^18^, essential for astrocyte infiltration into the γ axon bundle. The *orion* gene encodes two proteins: Orion-A and Orion-B, with functional redundancy in MB pruning ^18^. Moreover, secreted Orion recognizes the “eat-me” signal phosphatidylserine (PS) and acts as a molecular bridge between PS and the phagocytic receptor Drpr (*draper*), which is present in phagocytic epithelial cells during the clearance of *Drosophila* dendritic arborization (dda) neurons (Ji et al., 2023). We have recently demonstrated that Drpr also serves as the glial receptor for Orion during MB remodeling ^19^. Interestingly, in brain regions where glial infiltration is not required for neuronal remodeling (because glia is already present prior to pruning, such as around Corazonin or PDF-Tri neurons in the ventral nerve cord) Orion is nonetheless required for phagocytosis ^20^. Thus, Orion can induce both glial infiltration and phagocytosis. In MBs, γ neuron pruning requires the coordinated execution of glial infiltration and phagocytic clearance, likely involving activation of the same receptor, Drpr. How a single ligand as Orion can elicit either glial infiltration or phagocytosis, and how these two functions are mechanistically decoupled, remain fundamental unresolved questions.

Ligand presentation, including its spatial organization, density and mechanical context, has been proposed as a key determinant of receptor engagement and downstream signaling ^21, 22^. Overall, glycosaminoglycans (GAGs) play a central role in this process by concentrating and clustering ligands for presentation at the cell surface ^23, 24^. In addition, GAG-associated ligands can be internalized via heparan sulfate proteoglycan (HSPG)-mediated endocytosis and routed to endosomal compartments, including pathways involved in exosome biogenesis and trafficking ^25, 26^. Moreover, the progressive accumulation of ligand on GAGs over time, might generate a dynamic reservoir of both free and bound ligand, potentially expanding their functional role ^27^. Orion contains putative GAG-binding consensus sequences required for neuronal remodeling ^18^, suggesting that GAGs may serve as γ neuron-associated presenters of Orion to glial cells.

Here we sought to determine whether glial responses during neuronal pruning depend on the mode of ligand presentation. We demonstrated a biochemical association between Orion and heparan sulfate (HS) proteoglycans and show that HSs are required for γ axon pruning. Furthermore, tethering Orion to the γ neuron plasma membrane, mimicking its interaction with neuronal proteoglycans-associated GAGs, is sufficient to induce glial infiltration into γ axon bundles. However, this association fails to induce phagosome acidification, which instead requires both, a GAG associated Orion and most probably a mobile, “free” pool of Orion. Finally, we identified exosomes carrying GAG-bound Orion as potential contributors to neuronal remodeling. Together, our findings provide the first evidence that glial infiltration can be uncoupled from phagocytosis. Furthermore, our data indicate that the mode of ligand presentation critically influences glial responses and highlight a central role for GAGs in modulating Orion availability to differentially regulate astrocyte infiltration and phagocytosis and consequently, the efficiency of axon pruning and debris clearance.

## Results

### Purified Orion binds to heparan sulfate proteoglycans

We previously demonstrated that Orion binds to γ axon surfaces, accumulating at the tips of the axon lobes. Based on sequence analysis, three putative GAG-binding motifs were identified and mutation of any of them failed to rescue the unpruned axon phenotype in an *orion* mutant background, thus suggesting that this process was GAG dependent and could require direct interaction between Orion and GAGs ^18^.

In order to determine the implication of GAGs in MB pruning, we first wished to recombinantly produce Orion and biochemically assess its degree of association to GAGs as well as the nature of the GAG to which it binds. Although we expressed Orion using a homologous system (insect cells using the baculovirus expression system (Supplementary Fig. 1) to favour proper folding, potential native posttranslational modifications and secretion in the cell medium, expression yields were constantly very low (100-500 µg/L of medium), despite complete infection. Mammalian (HEK) or bacterial (*E. Coli*) expression systems did not give rise to better soluble yields (data not shown). Association of Orion to GAGs was first determined using a solid-phase assay that mimics, to some extent, the cell membrane-anchored GAGs presentation. For that purpose, reducing end biotinylated HS was immobilized on top of a streptavidin-coated sensorchip, and surface plasmon resonance (SPR) monitoring was used to measure changes in refractive index caused by the interaction when the protein samples were flowed across the immobilized HS surface. Binding responses were observed for each Orion concentrations and the experimental binding curves were best fitted to a bivalent analyte model (Fig. 1A), in agreement with the existence of distinct HS binding sites (see below). Kinetic analysis of the binding curves returns ka1 and kd1 values of 2.35 x 10^4^ M^-1^.s^-1^ and 1.19 x 10^-2^ s^-1^ respectively thus an affinity, for the first binding event, of KD1 = kd1/ ka1 of 506 nM. Such value should be considered as an approximative “apparent affinity” given the complexity of the system (several possible binding sites on the protein and, as it is common with HS, the existence of distinct oligosaccharidique sequences along the GAG chain, theoretically featuring distinct binding capacities). Competition assay, whereby Orion was preincubated with different GAGs before its injection over the HS sensorchip, showed that various chondroitin sulfate types (A, D, and E) were only able to partially inhibit Orion/HS interaction, while HS used as a competitor fully abrogated the interaction (Supplementary Fig. 2), demonstrating that HS is the principal GAG ligand of Orion.

**Figure 1.**
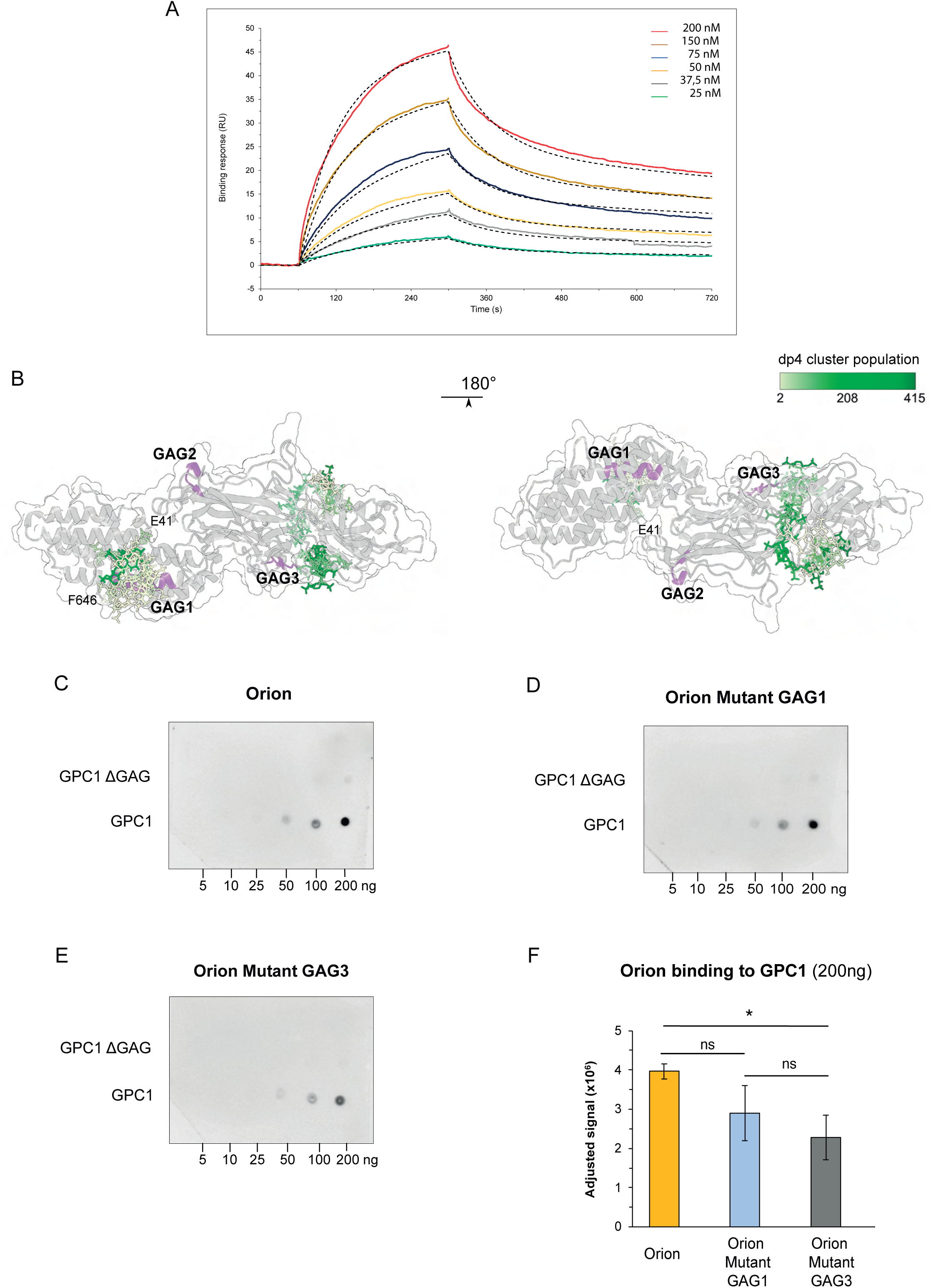
Orion binds to HS. (A) Orion was injected during 4 min over a heparan sulfate-activated sensorchip at a flow rate of 30 μL/ml after which the surface was washed for 8 min with running buffer. The binding response in resonance units (RU) was recorded as a function of time for Orion at (from top to bottom): 200, 150, 75, 50, 37.5 and 25 nM (coloured lines) and fitted to a bivalent binding model (dotted lines). (B) Visualization of the docking poses of an heparan sulfate dp4 (IdoA2S-GlcNS6S)2 on Orion AlphaFold3 model (residues 41-646). The dp4 are represented in sticks, coloured according to the number of members in each cluster. Predicted GAG binding domains are coloured in purple. (C-E) The binding of Orion wild type, Orion Mutant GAG1 and Orion Mutant GAG3 proteins to Glypican-1 (GPC1) was assayed by overlay dot blot. Increasing quantities of GPC1 and its mutant lacking all its GAG attachment sites (ΔGAG) were spotted on a nitrocellulose membrane before incubation with 3 µg of each Orion protein. Bound Orion was revealed using a horseradish peroxidase-coupled anti-His antibody and chemiluminescence detection. (F) Overlay dot blot statistics (n=3).

In order to get insight into the respective importance of the HS binding sites, we first used AlphaFold3 to predict Orion tridimensional structure, and then calculate its electrostatic potential surface (Supplementary Fig. 3). Orion model shows an elongated shape organized in two main regions, the N-terminal domain (41-256), composed of 7 α-helices, and a C-terminal domain mostly composed of β-sheets (Supplementary Fig. 3A). Basic amino acid sequences, identified as putative GAG binding sites are solvent exposed and found within an α-helix of the N-terminal domain (GAG1) and within the C-terminal domain (GAG2 and 3). Interestingly, GAG1 and GAG3 were located in large positively charged regions (blue colours), while GAG2 site positioned in a region with lower electrostatic potential. We next docked an HS tetrasaccharide (dp4) on the Orion tridimensional model (Fig. 1B). We observed a close contact between dp4 and GAG1 and GAG3 sites, involving several typical GAG binding residues (Supplementary Fig. 3B). In agreement with the electrostatic analysis of the Orion surface no dp4 docked onto the GAG2 sequence. Taken together, these results suggest a possible involvement of the GAG1 and GAG3 sequences in GAG recognition and prompted us to experimentally determine their respective importance in HS interaction.

Orion Mutant GAG1 and Orion Mutant GAG3 proteins were thus expressed and purified with the same strategy applied for the wild type protein (Supplementary Fig. 4). GAG2 mutant expression/secretion was too low to be purified successfully, suggesting a possible role of this sequence in protein folding (data not shown). Since glypicans are major neuronal glycoproteins harbouring HS chains, we sought firstly, to determine whether Orion binds to glypican-1 (GPC1) through its HS chains and secondly, to establish the effect of the GAG binding domains GAG1 and GAG3 on its binding.

For this, GPC1 and a mutant lacking the GAG attachment sites (GPC1 ΔGAG, S486A, S488A, S490A) were spotted onto a nitrocellulose membrane and incubated with 3 µg of Orion wild type, Orion Mutant GAG1 or Orion Mutant GAG3. Strong binding of Orion was observed with GPC1, whereas no binding was detected with the ΔGAG mutant (Fig. 1 C-F), indicating that Orion interacts with the HS chains associated with this proteoglycan. Comparison of the binding capacity of Orion wild type, Orion Mutant GAG1 and Orion Mutant GAG3 showed that Orion Mutant GAG3 only features a significantly reduced binding to GPC1 (Fig. 1 D, F), suggesting that the GAG3 site plays a key role in mediating Orion binding to HS.

Taken together, these results show a direct interaction between Orion and HS and strongly support the existence of different HS binding regions in Orion. Although binding is significantly reduced in GAG3 mutant compared to wild type, residual interaction with HS is still observed, suggesting that additional binding may occur, potentially through GAG1 sequence or other non-identified residues.

Since mutations in GAG3 binding domain of Orion gives the lowest percentage of rescue of the Orion mutant compared to the GAG1 domain ^18^ and because the Orion Mutant GAG3 protein exhibits the strongest lost in binding to GPC GAG chains, we focused our subsequent *in vivo* analysis on the GAG3 motif.

### Binding of Orion to GAGs is required for glia infiltration during neuronal remodeling

To determine at which step of axonal pruning the GAG3 motif contained in Orion is required we explored its role in glia infiltration into the MB γ axon bundle. The role of glia infiltration in MB pruning has been already demonstrated ^13, 35, 16^, as well as the key role of Orion in this infiltration. We visualized MBs by a membrane-targeted GFP (*UAS-CD8-GFP*) under the control of *201Y-GAL4* and co-stained glial cells with an antibody against Drpr at 6 h after pupa formation (APF). Glial extensions infiltrate γ axon bundles in wild type (Fig. 2 A-A”, E), opposite to *orion*^1^ (an *orion^null^* created by EMS mutagenesis ^18^) in which glia infiltration does not occur at 6 h APF (Fig. 2 B-B”, E). Expression of an *UAS-orion* construct driven by the γ neuron driver *201Y-GAL4* in an *orion*^1^ context rescued glia infiltration into the γ bundle (Fig. 2 C-C”, E). Interestingly, expression of a construct containing mutations in the GAG3 Orion binding domain (*UAS-orion-Mut GAG3*) in an *orion*^1^ context did not rescue glia infiltration compared to wild type (Fig. 2 D-D”, E). We performed similar experiments in *orion*^Δ*C*^ (an *orion* mutant created by the CRISPER-Cas9 system ^18^) and, as before, glial extensions did not infiltrate the γ bundle compared to a wild type (Supplementary Fig. 5).

**Figure 2.**
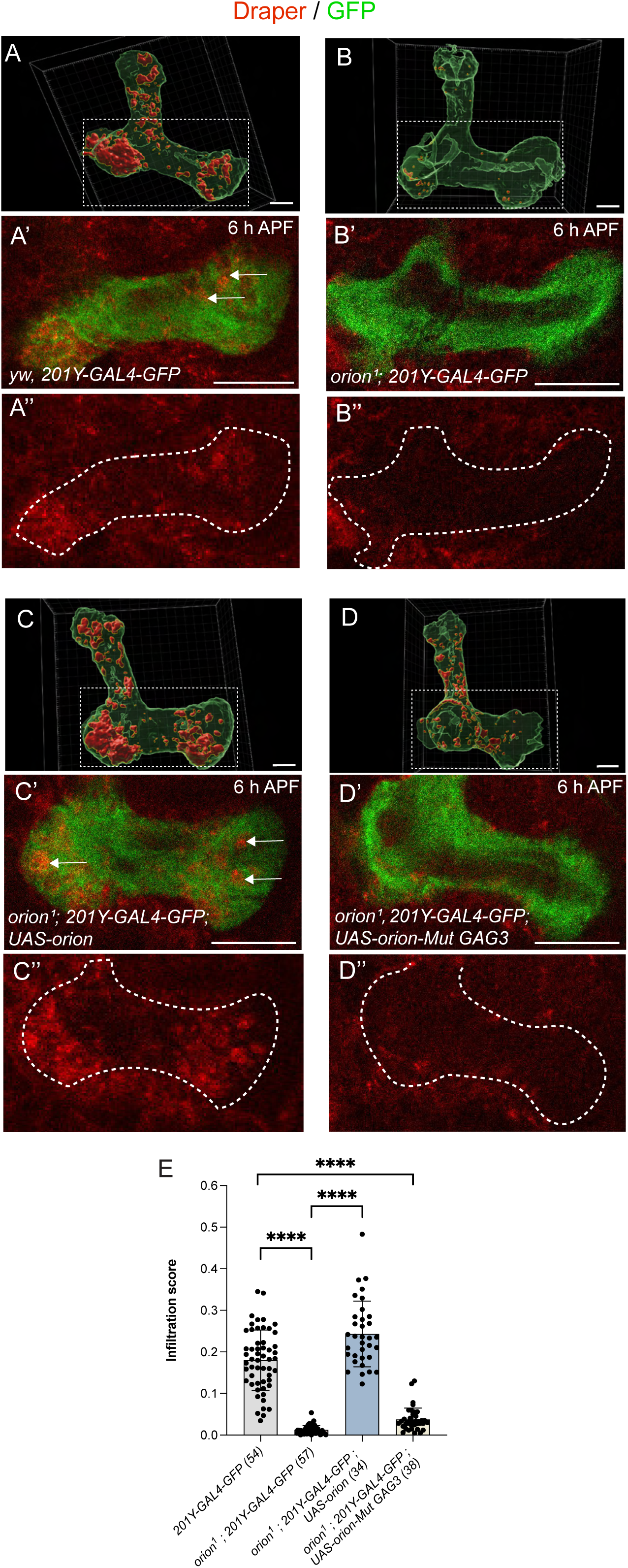
The association of Orion with GAGs is required for astrocytic infiltration into MB γ axon bundles at the onset of pruning. (A-D) Astrocytic infiltration into the γ axon bundle was analysed using the Imaris software, where the γ axon surface was demarcated in green and the astrocytic processes inside the axon in red. (A’-D’) Single confocal planes of 6 h APF brains expressing *201Y-GAL4*-driven *UAS-mCD8-GFP* (green). Anti-Drpr antibody (red) was used to visualize astrocytic processes. (A”-D”) Anti-Drpr antibody staining (red) in which the regions corresponding to the medial axon lobes were delimitated with discontinuous white lines. Experiments were performed in wild type (A panels), *orion*^1^ (B panels), *orion*^1^ expressing *201Y-GAL4*-driven *UAS-orion* (C panels) and *orion*^1^ expressing *201Y-GAL4*-driven *UAS-orion-Mut GAG3* (D panels). Astrocyte extension infiltration was labelled by arrows in A’ and C’. (E) Quantification of the γ lobe infiltration score by astrocytes (dorsal and medial lobe). Genotypes are listed in the supplementary Material and Methods (List of fly strains). Results are mean ± s.e.m. *n* values are indicated in a parenthesis for each condition. Replicated three times. \*\*\*\**P*<0.0001 (Mann–Whitney *U*-test). Scale bars: 10 µm, A-D and 20 µm, A’-D’.

We validated that the protein encoded by *Myc*-tagged *Orion-Mut GAG3* was normally secreted outside of neurons. For this, we analysed its presence at the “hole” like structures contained in the MB γ bundle at 6 h APF which correspond to vestiges of disappeared varicosities ^13^. If a protein is secreted by γ neurons, we expected it to occupy the inside of the holes. Accordingly, we found that Orion-Mut GAG3 occupied the inside of holes, in both a wild type and an *orion^-/-^*context, in which hole-like structures were smaller most probably due to the absence of glia infiltration (Supplementary Fig. 6 A-A”, C-C”). These results differed from the one observed when the transmembrane receptor Drl, *UAS-drl,* used as a control for the absence of secretion, was driven in γ neurons by *201Y-GAL4*. In this case, expression of Drl was limited to scattered dots randomly distributed in the periphery of the holes (Supplementary Fig. 6 B-B”). Taken together, our data strongly suggest that the Orion GAG3 binding domain is required for glial infiltration and point to GAGs as essential molecules required for MB γ neuron presentation of Orion to surrounding astrocytes.

### Dally and Dally-like glycoproteins localize with Orion and are essential for neuronal remodeling

Since Orion binds to HS chains *in vitro* (including those on GPC1), we investigated the association of Orion to *Drosophila* proteoglycans Dally (homologue of vertebrate GPC1) and Dally-like protein (Dlp) which are also associated to HS chains and are both highly expressed in the *Drosophila* nervous system ^36^. We took advantage of the fact that the fine temporal transcriptional landscape of MB γ neurons was recently described and a corresponding resource is freely accessible: https://www.weizmann.ac.il/mcb/schuldiner/resources/expression-profile to determine the mRNA developmental expression of these proteoglycans. Noteworthy, *dally* follows a pattern of expression very similar to *orion* (named also *l(1)G0193*), ^18^ (Supplementary Fig. 7 A), being highly transcribed at 3 h APF and dramatically decreasing at 9 h APF (Supplementary Fig. 7 B). Furthermore, *dlp* was highly transcribed in a similar temporal window, with a peak of expression at 6 h APF (Supplementary Fig. 7 C). The concomitant temporal expression of Orion with Dally and Dlp in γ neurons is consistent with a potential interaction between Orion and these proteoglycans. Moreover, to investigate this possibility, we expressed *UAS-dally-GFP* (Fig. 3 A-A”, C) and *UAS-dlp-GFP* (Fig. 3 B-B”, D) in MBs using *201Y-GAL4*, and analysed their colocalization with Myc-tagged *UAS-orion*. We observed 40.9% of colocalization between Orion and Dally and 34.7% between Orion and Dlp. These results support the hypothesis that Orion associates with proteoglycans in γ neurons and raise the possibility that Dally and Dlp contribute to presenting Orion to glial cells.

**Figure 3.**
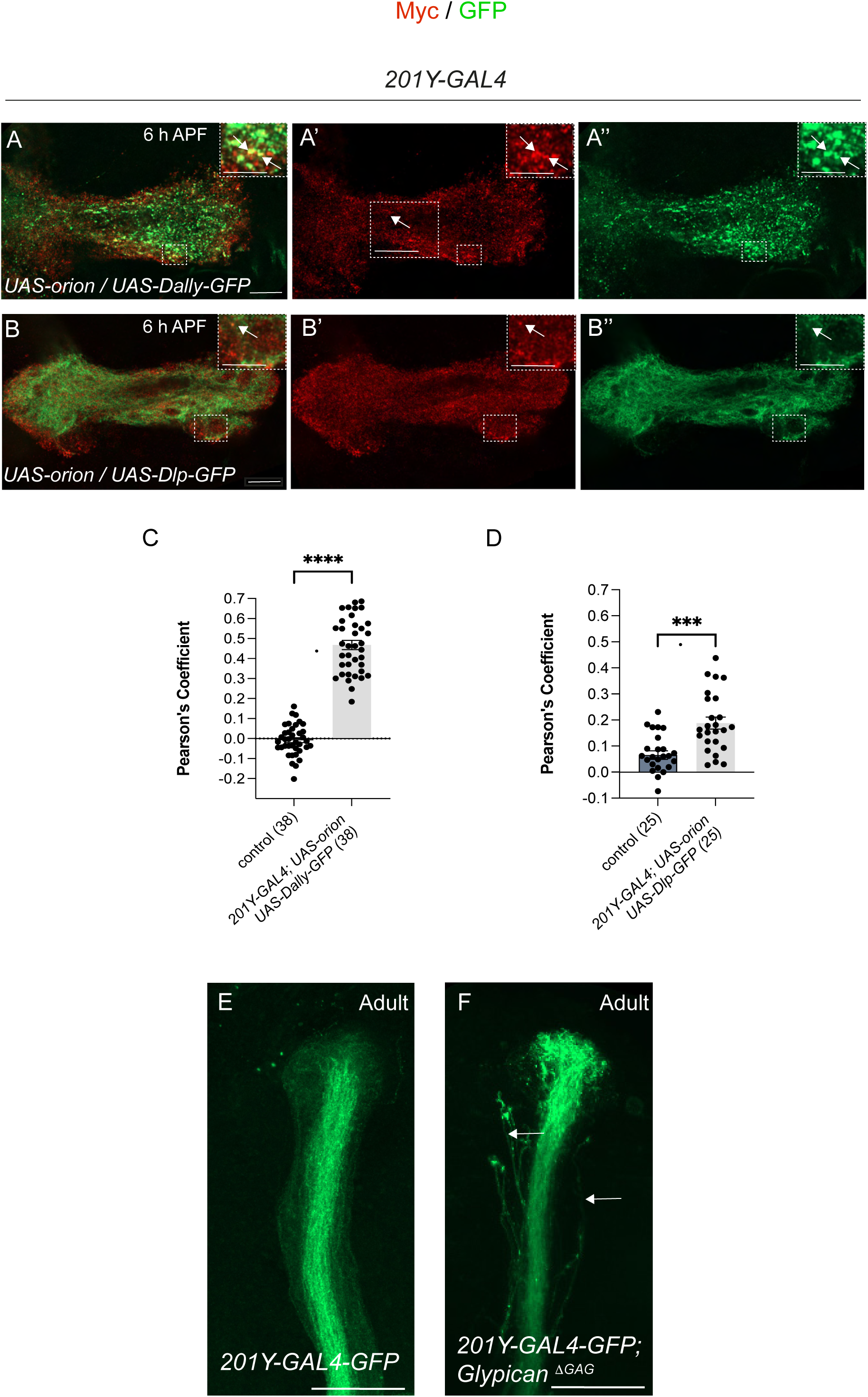
The association of Orion with heparan sulfate proteoglycans is required for MB remodeling. (A-A’’; B-B”) Single confocal planes of 6 h APF brains expressing *201Y-GAL4*-driven *UAS-orion* (Myc tagged) and *UAS-dally-GFP* (A-A’’) or *UAS-dlp-GFP* (B-B”) were used to analyze colocalization levels. Orion was visualized with an anti-Myc antibody (red) and Dally and Dlp were GFP tagged (green). Insets show high-magnification planes of the regions enclosed by a discontinued line in each image. The Pearson correlation coefficient was shown in C, D graphs and was used to quantify the degree of colocalization between Orion-Myc and Dally-GFP (C) and between Orion-Myc and Dlp-GFP (D). Control values in the graphs were obtained from the same quantified surface area for Dally-GFP or Dlp-GFP, respectively, after a 180° rotation to estimate the level of random colocalization. Confocal Z-stacks of adult brains expressing *201Y-GAL4*-driven *UAS-mCD8-GFP* in wild type (E) and *Glypican^ΔGAG^* (F). Individual unpruned γ axons were indicated by arrows in F. Genotypes are listed in the supplementary Material and Methods (List of fly strains). Results are mean ± s.e.m. *n* values are indicated in a parenthesis for each condition. Replicated three times. \*\*\*\**P*<0.0001 and \*\*\**P*<0.001 (Mann–Whitney *U*-test). Scale bars: 5 µm (A-A’’; B-B”) and 20 µm E, F.

Finally, to assess involvement of GAGs in pruning we analysed pruning phenotypes in adult flies containing double mutations in Dally and Dlp GAG attachment sites using the *Drosophila* line Glypican ^λGAG^ ^37^. Visualization of the corresponding MBs using *UAS-GFP* expression driven by *201Y-GAL4* showed the presence of unpruned γ axons in 100% of MBs observed (n ≥ 10 MBs) compared to controls, in which MBs displaying unpruned axons were not observed (0% MBs with unpruned axons, n ≥ 10 MBs) (Fig. 3 E, F).

Together these results point to HSPGs as major neuronal anchors of Orion to induce astrocyte infiltration.

### Orion tethered to γ neuron plasma membrane is sufficient for glia infiltration into MB γ neuron at the onset of metamorphosis

As Dally and Dlp are linked to the plasma membrane, we hypothesized that tethering Orion to plasma membrane with a CD2 transmembrane domain (Orion-CD2) could mimic the effect of the Orion association to GAGs. In addition, it was already shown that Orion-CD2 ^38, 19^ was functional and able to bind the Orion receptor Drpr. Consequently, we expressed in *orion*^1^ mutant MBs two different constructs: *UAS-orion-CD2* and *UAS-orion-CD2-myc* under the control of *201Y-GAL4.* Opposite to *orion*^1^ MBs which lack glia infiltration (Fig. 4 A-A’, E), we observed a very high level of glia infiltration with both constructs at 6 h APF (Fig. 4 B-C panels, E). We also performed a similar experiment in an *orion^ΔC^* mutant background and a high level of glia infiltration was also observed in *orion^ΔC^*; *UAS-orion-CD2* flies compared to *orion^ΔC^*alone (Supplementary Fig. 8).

**Figure 4.**
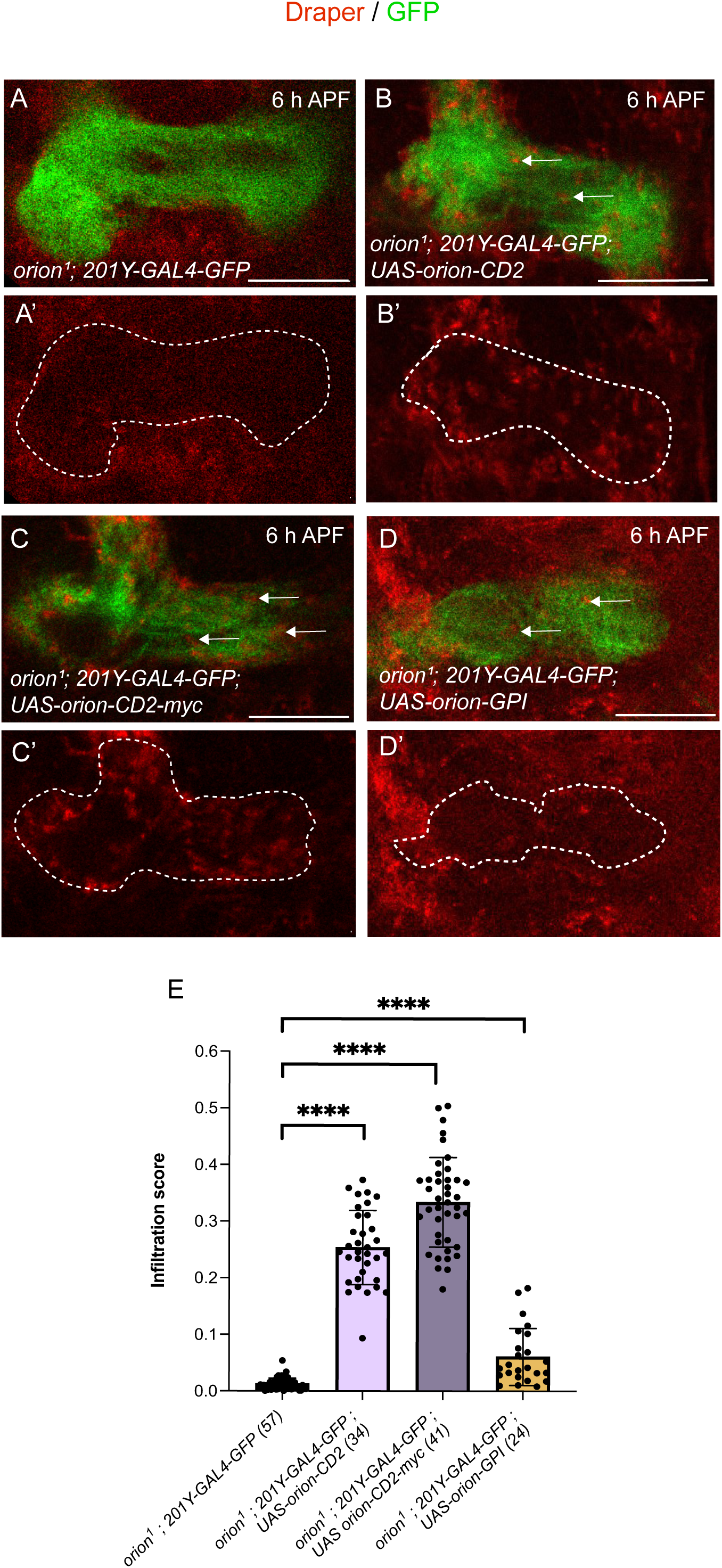
The association of Orion to plasma membrane is sufficient for astrocytic infiltration into MB γ axon bundles at the onset of pruning. (A-D) Single confocal plans of 6 h APF brains expressing *201Y-GAL4*-driven *UAS-mCD8-GFP* (green). Anti-Drpr antibody (red) was used to visualize astrocytic processes. (A’-D’) Anti-Drpr antibody staining (red) in which the regions corresponding to the medial axon lobes were delimitated with discontinuous white lines. Experiments were performed in *orion*^1^ (A panels), *orion*^1^ expressing *201Y-GAL4*-driven *UAS-orion-CD2* (B panels), *orion*^1^ expressing *201Y-GAL4*-driven *UAS-orion-CD2-myc* (C panels) and *orion*^1^ expressing *201Y-GAL4*-driven *UAS-orion-GPI* (D panels). Astrocyte extension infiltration was labelled by arrows in B-D. (E) Quantification of the γ lobe infiltration score. Genotypes are listed in the supplementary Material and Methods (List of fly strains). Results are mean ± s.e.m. *n* values are indicated in a parenthesis for each condition. Replicated three times. \*\*\*\**P*<0.0001 (Mann–Whitney *U*-test). Scale bar: 20 µm.

In a second set of experiments, we associated Orion with the plasma membrane of γ neurons using a glycosyl-phosphatidyl-inositol (GPI) sequence and expressed *201Y-GAL4*-driven *UAS-orion-GPI* in MBs in an *orion*^1^ mutant background. We observed a statistically significant increase in glial infiltration in these flies, compared with *orion*^1^ mutants alone, although the level of infiltration was lower than that obtained with *UAS-orion-CD2* (Fig. 4 D-D’, E). As a control, we verified that Orion-CD2; Orion-CD2-Myc and Orion-GPI were all detected at the surface of the γ axon lobes (Supplementary Fig. 9). Together these results suggest that plasma membrane-tethered Orion is sufficient to promote glia infiltration in the γ neuron bundle.

### Tethering Orion to γ neuron plasma membrane abolishes phagocytosis

We next investigated if the presentation of Orion tethered to γ neuron plasma membrane was also sufficient to induce phagocytosis. Accordingly, we used the membrane associated pH sensor *UAS-MApHS* that allows to detect acidic structures in phagocytes after engulfment as a read out of phagocytosis ^39^.

We clearly evidenced the presence of phagolysosomes in γ axon bundles in controls expressing *201Y-GAL4* driven *UAS-MApHS* in MBs at 6 h APF (Fig. 5 A). Phagosomes appeared as spherical structures of diverse sizes and were distributed all along the MB dorsal and medial γ axon bundle. We focused on the MB medial bundle for quantifications and calculated the ratio between the area occupied by phagosomes and the total MB area. In controls we observed phagosomes occupied an area significantly higher to the one occupied by phagosomes in *orion^ΔC^*, which lack phagocytosis, (Fig. 5 A, B, G). Phagocytosis was rescued in *orion^ΔC^* expressing *201Y-GAL4* driven *UAS-orion* (Fig. 5 C, G). No phagocytosis was observed neither in *orion^ΔC^* expressing *201Y*-GAL4 driven *UAS-orion-Mut GAG3* nor in *orion^ΔC^*expressing *201Y-GAL4* driven *UAS-orion-CD2* or *UAS-orion-CD2-myc* (Fig. 5 D-G). To our knowledge, the finding that MB expression of Orion-CD2 induces glial infiltration while preventing phagocytosis represents the first evidence that glial infiltration and phagocytosis can be dissociated. Furthermore, these results suggest, first, that in the absence of glial infiltration (as observed in *orion-Mut GAG3*) phagosome acidification does not occur, and second, that glia infiltration alone is not sufficient to enable glia to phagocytose axonal material (as observed in *orion-CD2*). Thus, in the wild type situation, phagocytosis might require both, glial infiltration mediated by GAG-bound Orion and the presence of a free pool of Orion.

**Figure 5.**
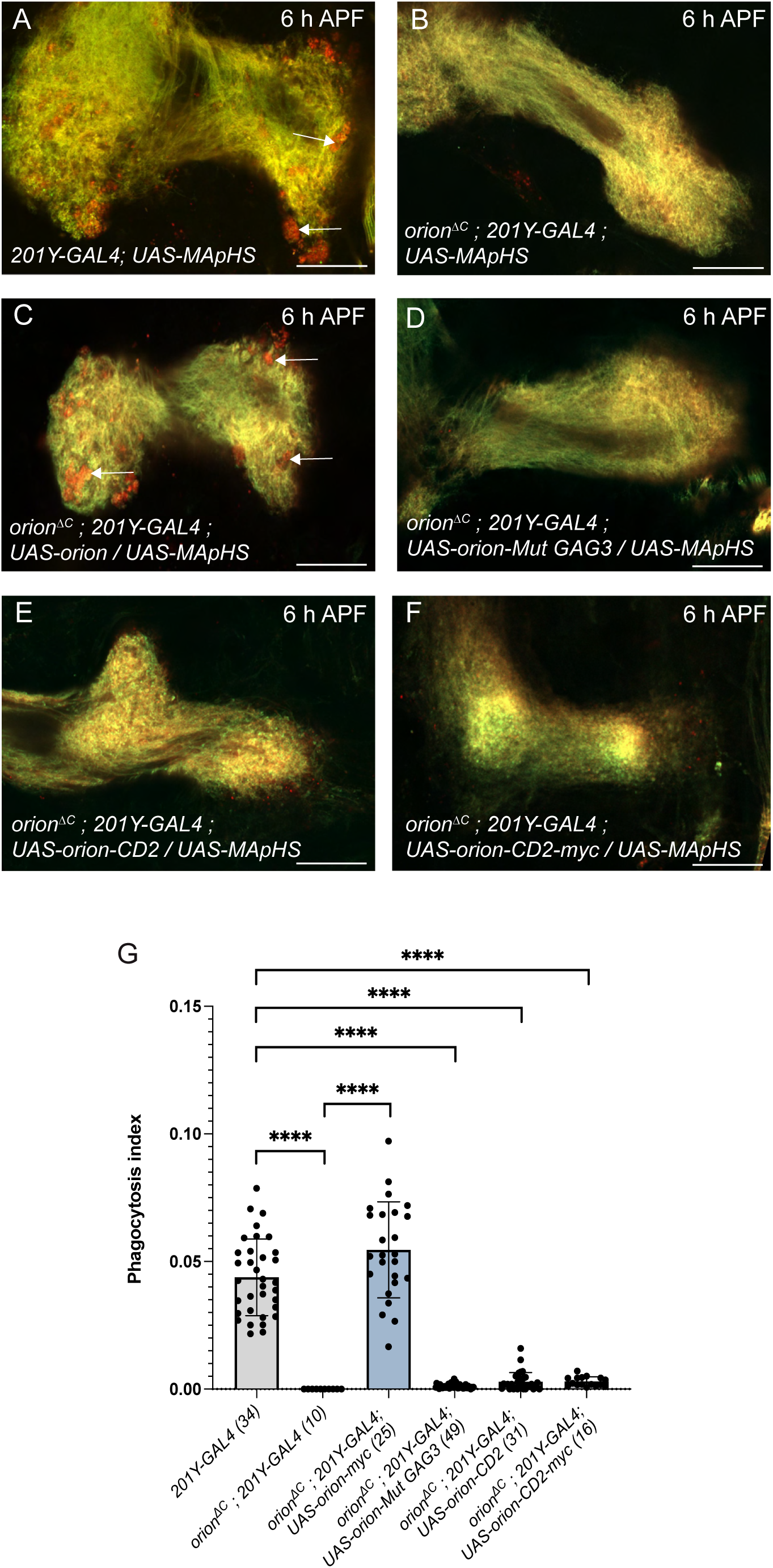
Orion tethered to plasma membrane fails to induce phagosome acidification in astrocytes. (A-F) Single confocal planes of 6 h APF brains expressing *201Y-GAL4*-driven *UAS-MApHS* (green/red). Green and red colours merge all along the γ axon bundle except in the regions in which axonal material is included inside glial acidic phagosomes leading to red inclusions (arrows in A and C). Phagosomes were observed in wild type (A) and *orion*^Δ*C*^ expressing *201Y-GAL4*-driven *UAS-orion* (C). In *orion*^Δ*C*^ expressing *201Y-GAL4*-alone (B) or in the presence of *201Y-GAL4*-driven: *UAS-orion-Mut GAG3* (D*), UAS-orion-CD2* (E), *and UAS-orion-CD2-myc* (F) phagosomes were absents or in very low amount. (G) Quantification of the phagocytosis index in the medial γ lobe. Genotypes are listed in the supplementary Material and Methods (List of fly strains). Results are mean ± s.e.m. *n* values are indicated in a parenthesis for each condition. Replicated three times. \*\*\*\**P*<0.0001 (Mann–Whitney *U*-test). Scale bar: 20 µm.

We recently showed the presence of a MB “headless” phenotype in 6 h APF pupae expressing *201Y-GAL4* driven *UAS-orion-CD2* in MBs ^19^, consisting of the loss of the top of the lobe at this time point. Since *UAS-orion-CD2* expression in MBs prevented phagocytosis, we anticipate that if part of the γ lobe was lost either some axonal material was engulfed by glia but not digested or phagosomes could be produced earlier in development in *UAS-orion-CD2* expressing MBs. For this reason, we explored the possibility that late phagosomes could appear at larval and 2 h APF stages. In *201Y-GAL4* driven *UAS-orion-myc* and *UAS-orion-CD2*, we observed that the surface of phagosomes was higher at 2 h APF than at L3 (Supplementary Fig. 10). Curiously, the surface of phagosomes at 2 h APF was higher in *UAS-orion-CD2* and *UAS-orion-CD2 myc* individuals than in *UAS-orion* wild type; but drastically decreased at 6 h APF. Nevertheless, the phagosome surface observed in *UAS-orion-CD2* expressing MBs was low compared to the one observed at 6 h APF in *201Y-GAL4* driven *UAS-orion* wild type. These results suggested that Orion-CD2-presenting γ axons might initiate phagocytosis, however, due to the defective processing, the mechanism is aborted prematurely.

### Plasma membrane-tethered Orion in combination with partially GAG-bound Orion rescues γ axon phagocytosis and leads to adult MB pruning

The observation that plasma membrane-associated Orion is sufficient to induce glial infiltration but not normal phagocytosis, led us to hypothesize that efficient phagocytosis required a “free” (not bound to γ neuron plasma membrane) form of Orion to proceed. Thus, we tried to reconstitute the normal function of Orion in both infiltration and phagocytosis by combining two forms of Orion: one able to induce infiltration and a second one able to induce phagocytosis. We decided to take advantage of the fact that Orion-Mut GAG3 behaves as an Orion form with low ability to associate to GAGs (Fig. 1 E and Supplementary Fig. 6) and would consequently rescue phagocytosis in Orion-CD2-expressing γ axons without affecting glia infiltration. We first confirmed that glia infiltration was not compromised in *orion* mutants expressing of *201Y-GAL4*-driven *UAS-orion-CD2* combined with *UAS-orion-Mut GAG3* (Supplementary Fig. 11 A-C). Moreover, we observed that concomitant expression of *201Y-GAL4* driven *UAS-orion-CD2* and *UAS-orion-Mut GAG3* in *orion^ΔC^* background led to a statistically significant increase of the phagocytosis area fraction compared to *201Y-GAL4* driven *UAS-orion-CD2* alone (Fig. 6 A-C).

**Figure 6.**
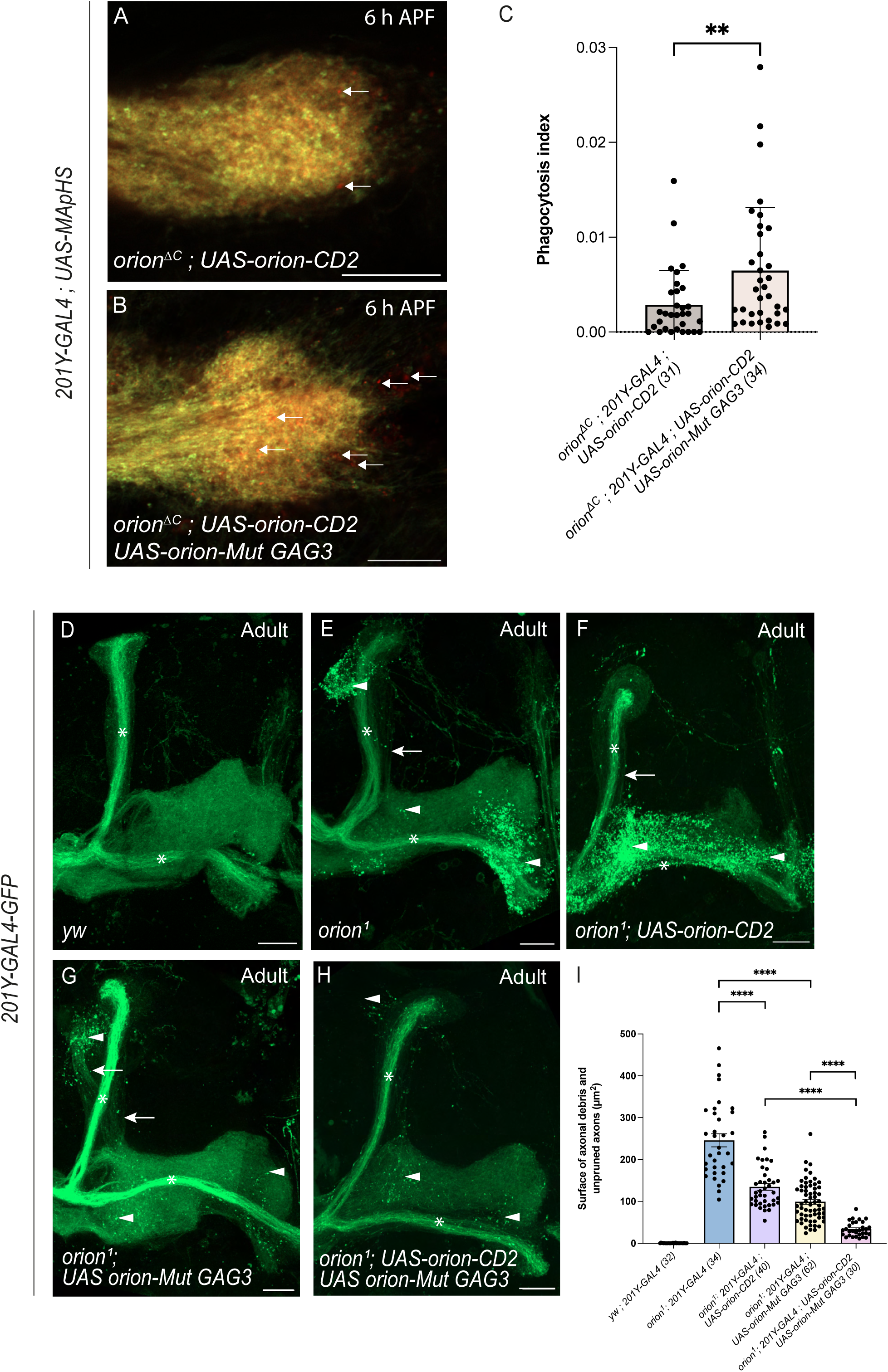
Orion associated to plasma membrane in combination with Orion unbound to GAGs *via* its GAG3 domain rescues phagocytosis and leads to adult MB pruning. (A, B) Single confocal plans of 6 h APF *orion*^Δ*C*^ brains expressing *201Y-GAL4*-driven *UAS-MApHS* (green/red) and *UAS-orion-CD2* (A) *or 201Y-GAL4*-driven *UAS-orion-CD2, UAS-orion-Mut GAG3* (B). Phagosomes are labelled by arrows. Note the high number of phagosomes present in B compared to A. (C) Quantification of the phagocytosis index in the medial γ lobe. (D-H) Confocal Z-stacks of adult MBs expressing *201Y-GAL4*-driven *UAS-mCD8-GFP* (green) in wild type (D), *orion*^1^ (E), *orion*^1^; *UAS-orion-CD2* (F), *orion*^1^; *UAS-orion-Mut GAG3* (G) and *orion*^1^; *UAS-orion-CD2 UAS-orion-Mut GAG3* (H). (I) Quantification of axonal debris and unpruned axon surface (μm^2^) in a whole MB. Genotypes are listed in the supplementary Material and Methods (List of fly strains). Results are mean ± s.e.m. *n* values are indicated in a parenthesis for each condition. Replicated three times. \*\*\*\**P*<0.0001 (Mann–Whitney *U*-test). Scale bars: 10 µm (A, B) and 20 µm (D-H).

Furthermore, we reasoned that the increase in the phagocytosis area fraction obtained in *201Y-GAL4* driven *UAS-orion-CD2* in combination with *UAS-orion-Mut GAG3* in *orion^ΔC^*background should result in increased adult γ axon pruning compared to *201Y-GAL4* driven *UAS-orion-CD2* alone. Thus, we quantified the total surface of axonal debris and unpruned axons in wild type, *orion*^1^, *orion*^1^ expressing either *201Y-GAL4* driven *UAS-orion-CD2* or *UAS-orion-Mut GAG3* alone and finally in *orion*^1^ expressing a combination of both *201Y-GAL4* driven *UAS-orion-CD2* and *UAS-orion-Mut GAG3* (Fig. 6 D-I). Interestingly, we observed that a combination of both *201Y-GAL4* driven *UAS-orion-CD2* and *UAS-orion-Mut GAG3* resulted in a rescue of the axonal debris and unpruned axon surface compared to *UAS-orion-CD2* or *UAS-orion-Mut GAG3* alone. Similar results were obtained when we used *orion^ΔC^* instead of *orion*^1^ for our experiments (Supplementary Fig. 11 D-I).

These data indicate that the combined presence of membrane-tethered Orion, which enables glial infiltration, and freely diffusible or weakly plasma membrane-attached Orion, which alone is insufficient to promote infiltration, restores glial phagocytosis and, consequently, adult axon and debris elimination. Thus, these findings suggest that distinct pools of Orion are required for the progression of MB axonal pruning, potentially corresponding to a GAG-bound form and a “free” (or a relatively free) form.

### Orion-associated to exosomes might be involved in γ axon elimination at the onset of pruning

We next explored the possibility that other forms of Orion could be involved in γ axon elimination, based on the fact that Orion was reported in two independent works as a protein associated to EV isolated from S2 cell supernatant ^40^ and from larval haemolymph ^34^. We first confirmed that Orion was present in larva haemolymph. For that, we expressed *UAS-orion Myc or GFP*-tagged in the fat body using the fat body specific *Lpp-GAL4* line and we analysed the corresponding larva haemolymph. Using an anti-Myc antibody we detected in haemolymph the 90 KD band corresponding to Orion by Western blot (Fig. 7 A). Moreover, we isolated exosomes from these haemolymphs using both EV-purification columns (Fig. 7 A) or by ultracentrifugation steps (Supplementary Fig. 12 A). Interestingly, a faint band corresponding to Orion was observed in the exosome-containing fraction with both purification methods, confirming that Orion also travels *via* exosomes. Furthermore, we used this assay to examine if Orion-Mut GAG3 was associated to exosomes. In theory, exosome biogenesis starts with endosome formation. Because Orion binds to GAGs that can enter the endocytic pathway it may be found in exosomes. In contrast, Orion-Mut GAG3, which have reduced binding to GAGs, should not be present in exosomes. To test this hypothesis, we used larva expressing *Lpp-GAL4* driven *UAS-orion-Mut GAG3*, extracted larva haemolymph and purified exosomes. Using an anti-Myc antibody we detected the expected 90 KD band by Western blot (Fig. 7 B) corresponding to Orion-Mut GAG3 in haemolymph extracts but not in exosomal fractions. We verified the integrity of our extracts using an α-Tubulin isoform, known as being present in exosomes ^34^, as a positive control (Fig. 7 A-B). Accordingly, the presence of Orion-Mut GAG3 in haemolymph provides additional evidence that Orion-Mut GAG3 can exist as a “free” protein, not associated with the plasma membrane. Furthermore, these findings also suggest that the GAG3-binding motif of Orion is required for its entry into the exosome biogenesis pathway. Since the role of the exosome pathway in γ axon pruning has not been previously reported, we first examined the localization of exosomes within γ axon bundles using the exosome marker CD63. To this end, we expressed *UAS-CD63-mCherry* in MBs using the *201Y-GAL4* driver and observed a strong accumulation of the marker in MB “hole-like” structures. As these structures correspond to regions of neuron-glia interaction during pruning ^13^, the presence of exosomes in these areas suggests that they may play a role in the γ neuron pruning process (Fig. 7 C-C’). Next, we assessed whether exosomes could colocalize with Orion in MBs. To this end, we co-expressed *UAS-CD63-mCherry* and *UAS-orion-Myc* in MBs using the *201Y-GAL4* driver and analysed their colocalization. We observed a statistically significant colocalization between CD63 and Orion (Fig. 7 D-D’, F)., compared to an irrelevant *UAS* (*UAS-mito-GFP*) expressed in MBs (Supplementary Fig. 12 B-B’). Additionally, we analysed the colocalization of Orion with proteins acting at late stages of the exosome biogenesis pathway, namely *UAS-Rab11-GFP* (Fig. 7 E-E’, F), *UAS-Rab7-GFP* and *UAS-LAMP-GFP* (Fig. 7 F and Supplementary Fig. 12 C-D), by expressing these proteins in MBs under the control of *201Y-GAL4*. For all three markers, a statistically significant colocalization was observed in the γ lobes compared with the control *UAS-mito-GFP*. Altogether, these data support a model in which, following binding to GAGs and endocytosis, Orion enters the exosome biogenesis pathway in γ axons and associates with secreted exosomes to mediate neuron-glia communication during developmental MB pruning. Finally, we sought to decipher the contribution of the exosome pathway to γ axon pruning. To address this, we expressed dominant-negative (*DN*) forms of proteins acting at late stages of exosome biogenesis. Because endocytosis has previously been shown to be required for MB pruning to proceed ^41^, we aimed to avoid perturbing this pathway and instead specifically target late endosomal or multivesicular body transport. Accordingly, we expressed *UAS-Rab11^DN^*, *UAS-Rab35^DN^*, and *UAS-Rab39^DN^* in MBs using the *201Y-GAL4* driver (Fig. 7 G-K), and assessed whether blocking the function of these proteins would affect pruning. Our results showed normal γ axon remodeling in controls (n=32). However, disrupting the function of Rab11, Rab35 or Rab39 lead to γ neuron remodeling defects (87.5% n=80, 88.24 % n=51 and 54.35 % n=46, respectively), characterized by the presence of uncleared debris and unpruned γ axons, a phenotype reminiscent of that observed at the onset of metamorphosis when glial function is impaired. Moreover, these data suggest that the exosome pathway is involved in Orion-mediated MB pruning and may represent and additional source of Orion (Fig. 8).

**Figure 7.**
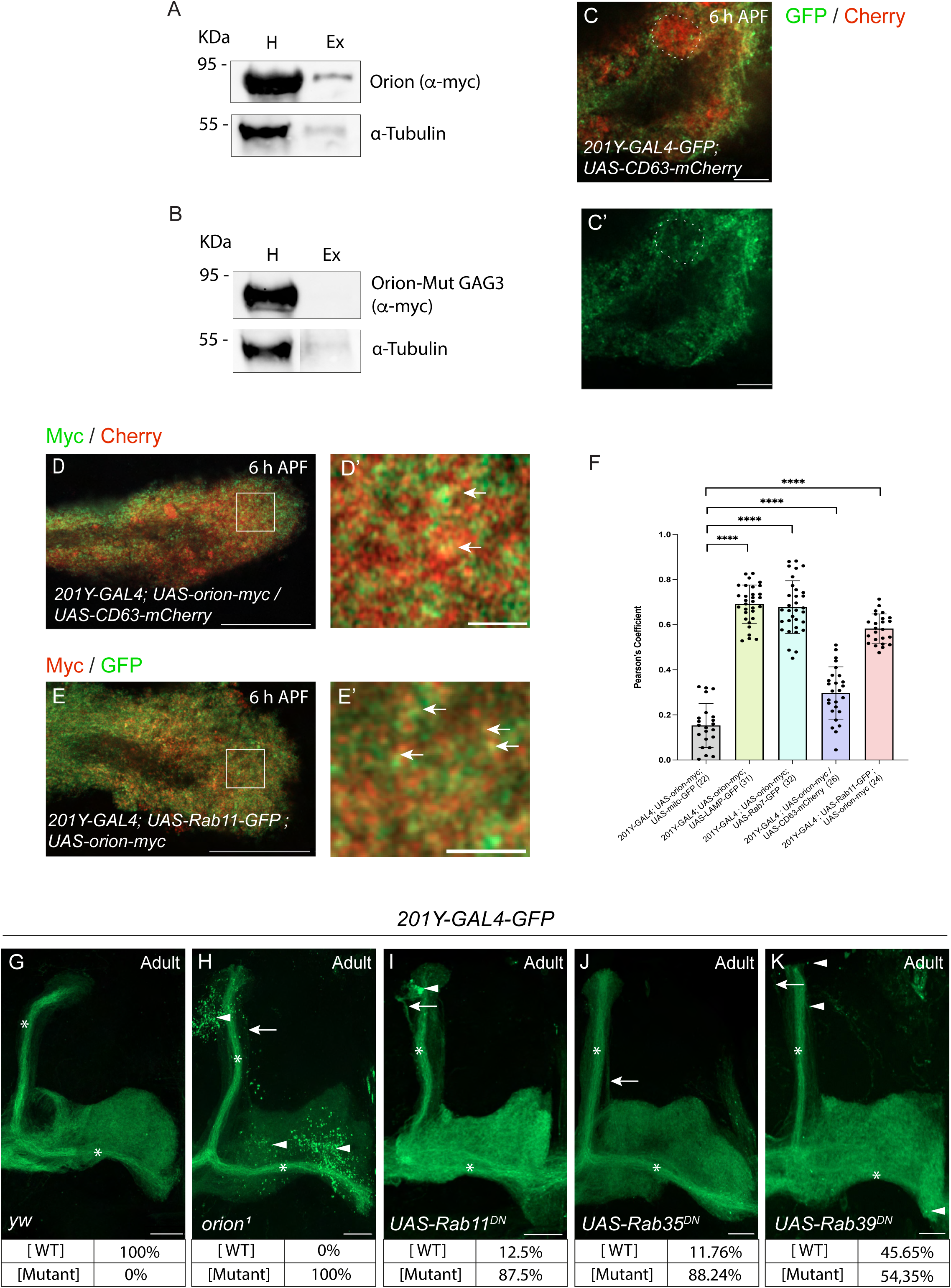
Exosome-carried GAG-bound Orion is required for γ neuron remodeling. (A, B) Immunoblot detection of Orion (A) and Orion-Mut GAG3 (B) in larval haemolymph extract (H lanes) and purified exosome fractions (Ex lanes) in *Lpp-GAL4-driven UAS-orion-myc* (A) and *Lpp-GAL4-driven UAS-orion-Mut GAG3* (B). α-Tubulin was used as an internal positive control. Anti-Myc and anti-Tubulin antibodies were used to detect Orion and α-Tubulin respectively. Note the presence of Orion in the exosome fraction, Ex lane in A. (C, C’) Single confocal plan of a 6 h APF brain expressing *UAS-mCD8-GFP* (green) *and UAS-CD63-mCherry* (red) driven by *201Y-GAL4*. A hole-like structure was delimitated with discontinuous white lines in the γ lobe (C). Note the high level of CD63 (red colour) present in this region. (D-E panels) Single confocal planes of 6 h APF brains expressing *201Y-GAL4-driven UAS-orion-myc* (green) and *UAS-CD63-mCherry* (red) (D-D’) or *201Y-GAL4-driven UAS-orion-myc* (red) and *UAS-Rab11-GFP* (green) (E-E’). D’, E’ are higher-magnification views from the corresponding square in D, E, respectively. Orion was visualized with an anti-Myc antibody (green in D and red in E). (F) The Pearson correlation coefficient, shown in the graph, was used to assess colocalization surface between Orion-Myc and several exosomal protein in the medial γ lobe: LAMP-GFP, Rab7-GFP, CD63-mCherry and Rab11GFP; compared with a random colocalization control between Orion-Myc and mito-GFP. (G-K) Confocal Z-stacks of adult MBs expressing *201Y-GAL4*-driven *UAS-mCD8-GFP* (green) in wild type (G), *orion*^1^ (H), *UAS-Rab11^DN^* (I), *UAS-Rab35 ^DN^* (J) and *UAS-Rab39 ^DN^* (K). Distribution of phenotypes: wild type (WT) or presence of unpruned axons and/or uncleared debris (Mutant), are respectively mentioned at the bottom of the pictures. Genotypes are listed in the supplementary Material and Methods (List of fly strains). Results are mean ± s.e.m. *n* values are indicated in a parenthesis for each condition in graph F. Replicated three times. \*\*\*\**P*<0.0001 (Mann–Whitney *U*-test). Scale bars: 5 µm (C, C), 20 µm (D, E, G-K) and 2 µm (D’, E’).

**Figure 8.**
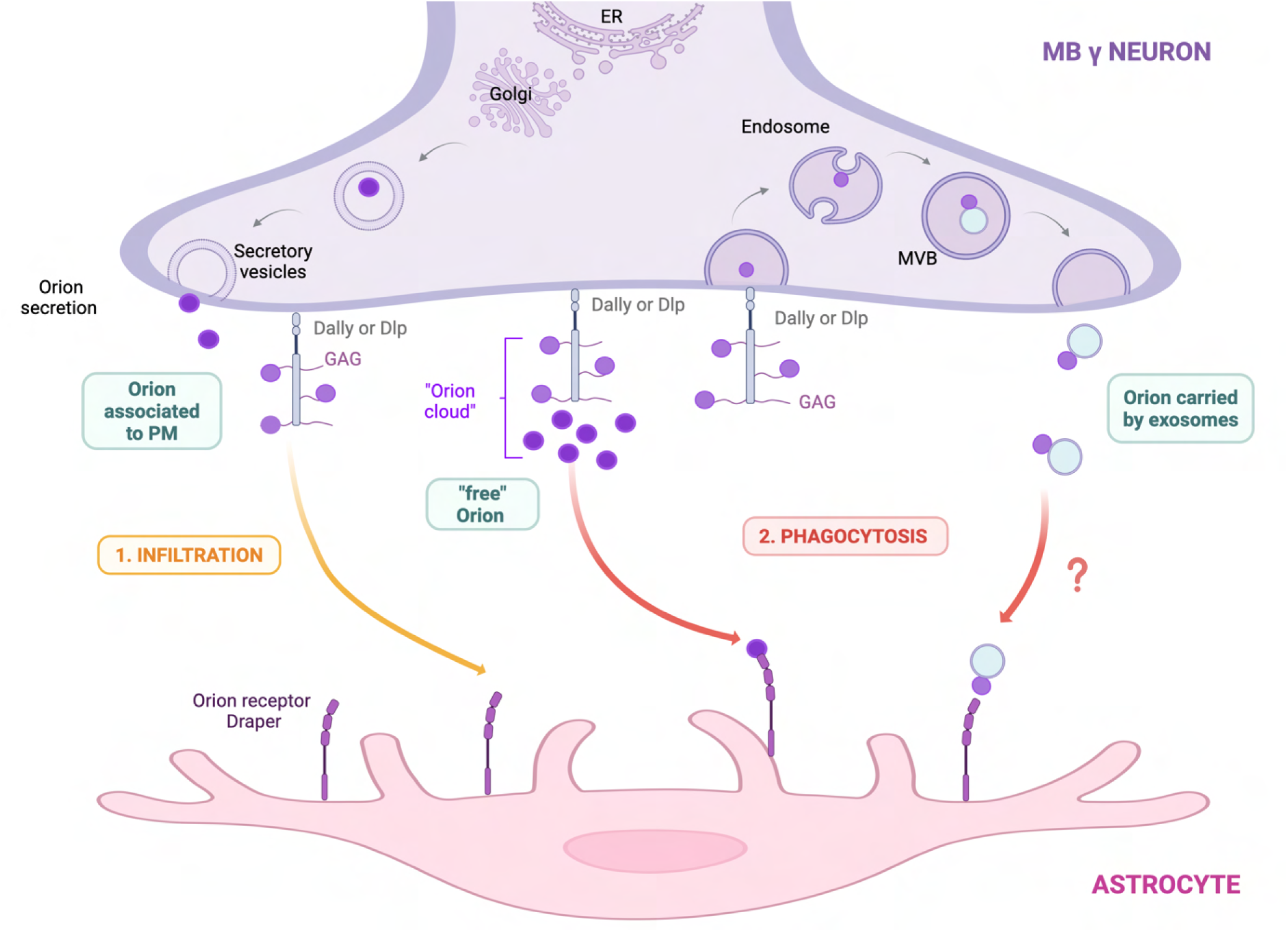
Model of Orion secretion, presentation, and astrocyte responses. Orion is secreted by γ neurons *via* the conventional Endoplasmic reticulum-Golgi pathway and released into the extracellular space through secretory vesicles. In the extracellular environment Orion binds GAGs, including those associated to Dally and Dlp proteoglycans, which present the ligand to surrounding astrocytes. This interaction triggers activation of the Drpr receptor and promotes astrocyte infiltration into γ axon bundles. Sustained Orion secretion may lead to its clustering on GAGs, generating a localized “ligand cloud” composed of both bound and “freely” diffusing Orion. This could establish a retained micro gradient that robustly activates the Drpr receptor and may drive appropriate phagocytosis of neuronal material by astrocytes. In addition, GAGs binding of Orion may engage the exosome-biosynthesis pathway in γ neurons. Orion-associated exosomes could serve as concentrated signalling platforms, further enhancing glial phagocytic responses. Abbreviations: MB: mushroom body, ER: endoplasmic reticulum, PM: plasma membrane, GAG: glycosaminoglycans, DLP: Dally-Like-Protein, MVB: Multivesicular body. This illustration was created with BioRender.com

## Discussion

Neuronal pruning requires the coordinated execution of glial infiltration and phagocytic clearance ^42, 7, 13, 18, 14, 15, 16, 35^, yet the mechanisms involved in the segregation of these steps *in vivo* remain unknown. *Drosophila* chemokine-like Orion has emerged as a key neuron-secreted ligand for glia infiltration into γ axon bundles and phagocytosis ^18, 20^. Both mechanisms seem mediated by the phagocytosis receptor Drpr (homologue to MEGF10 in mammals). Indeed, early L3-infiltrating glia displaying high levels of Drpr was observed when Orion was associated with plasma membrane ^19^ and phagocytosis of da neurons by phagocytic epithelial cells was demonstrated to depend on Drpr ^38^.

Our study directly examines how the neuronal presentation of a ligand to glia can modify glial responses depending on the way this ligand, in our case Orion, is presented. These differences may arise from the way Orion presentation modifies Drpr conformation, thereby activating and adapting specific signaling pathways. In accordance, differential receptor activation by membrane associate versus soluble ErbB1 ^43^ and EGF ligands was reported before ^44^. Here, we propose a model (Fig. 8) in which GAGs promote different modes of Orion presentation to astrocytes. Since Orion contains a signal peptide sequence allowing it to be secreted *via* a conventional protein secretion pathway, we propose that once in the extracellular space, Orion binds to GAGs associated with proteoglycans Dally and Dlp. In support to our model, we showed using biochemical approaches Orion direct binding to isolated HS chains, as well as HS associated with the glypican GPC1. In addition, we observed colocalization between the proteoglycans Dally and Dlp and Orion, as well as correlated expression of the three proteins in γ neurons early during metamorphosis. Together these data suggest that the Orion-glypican interaction has a functional role in pruning. In accordance, mutation in Dally and Dlp GAG attachment sites resulted in adult unpruned γ axons which might indicate that GAG deficiency on these proteoglycans prevents appropriate Orion presentation to glia and consequently adult axon pruning. Functionally, we mutated a GAG-binding site in Orion, the GAG3 domain, which exhibited the strongest biochemical interaction with GAGs among the three putative binding sites and is also required for MB pruning ^18^. We found that this interaction was necessary for glial infiltration, as mutations in the GAG3 binding site prevented glial infiltration into MB lobes. Furthermore, glial infiltration has been shown to depend on actin remodeling ^16^, which through Drpr activation might be regulated by the association between Orion and GAGs.

GAGs are well established regulators of ligand distribution and signaling. By binding secreted factors such as chemokines or morphogens, GAGs concentrate ligands at their site of production, prevent their diffusion, and promote the formation of localized gradients. In addition, GAGs act as scaffolds which present ligands to their receptors in a spatially controlled manner, thereby contributing to signaling specificity ^45, 46, 47, 48, 37^. In particular, HSPGs, including Dally and Dlp in flies, as well as Syndecans in vertebrates are well known regulators of secreted ligand signaling ^49^. Interestingly, Dally and Dlp have been shown to control the distribution and activity of multiple morphogens, including Dpp, Wingless and Hedgehog, by acting as co-receptors and regulators of ligand availability ^50^. In parallel, studies on chemokines have also demonstrated GAG binding stabilization of ligand gradients which enables their proper presentation to receptors ^51^. Importantly, GAG-binding of chemokines has been identified as essential for regulating chemotaxis, either in ensuring their correct positioning within tissues ^52^ or in directly regulating their signaling activities ^53^. Together, these findings support the idea that GAGs might actively shape Orion distribution and signaling output.

Because in MBs phagocytosis requires previous glia infiltration, we were unable to directly determine whether Orion-GAG binding was also required for phagocytosis in flies lacking glia infiltration. To address this, we mimicked the Orion-GAG interaction by tethering Orion to the γ-neuron plasma membrane using two different C-terminal tags: a CD2 (Orion-CD2) and a GPI (Orion-GPI). Although these constructs do not fully recapitulate the endogenous Orion-GAG interaction, we reasoned that they could help to determine whether membrane association of Orion is sufficient to induce glial infiltration, or whether other modes of GAG binding, for instance Orion association with the extracellular matrix, are required. Firstly, a high level of glia infiltration was observed in *orion* mutants expressing Orion on γ-neuron plasma membrane compared to *orion* mutant alone, which suggested that association of Orion to plasma membrane was sufficient to induce glia infiltration. Secondly, the increased level of glial infiltration observed in Orion-CD2 compared to Orion-GPI, suggested a requirement of a highly stable association between Orion and GAGs for glial infiltration. In accordance with this role of GAGs in MB-surrounding glia infiltration, it was recently proposed that a gradient of GAG-binding chemokines is first exposed on apoptotic cells to guide leukocytes to their target ^54^.

One striking result was that, in an *orion*^-/-^background, γ-neuron -expressed Orion-CD2 was sufficient to induce glial infiltration, but surprisingly phagocytosis was impaired, with only a very low amount of material being phagocytosed. Thus, effective phagocytosis might require either a form of Orion less tightly associated with the plasma membrane than Orion-CD2 or a form of “free” Orion in the extracellular space. Thus, when Orion-CD2 is presented to glia, it might lead to a suboptimal conformation of the Drpr receptor limiting activation of phagocytic pathways, which are typically mediated by Ced-6 ^55, 56^, Src, Shark ^57^ and associated signaling complexes Crk / Mbc (DOCK180) / ELMO / Rac1 ^58^. It was supported by the observation that co-expression of Orion tethered to plasma membrane with Orion mutated in its GAG3 binding motif in an *orion* mutant context rescued the phagocytic defect observed in Orion-CD2-expressing MBs. In addition, quantitative analysis of neuronal pruning revealed that these distinct Orion forms (Orion-CD2 and Orion-Mut GAG3) differentially orchestrated the timing and efficiency of adult γ neuron debris clearance *in vivo* (Fig. 6 I). Accordingly, we propose two functional pools of Orion coexisting during metamorphosis: a GAG-associated one which promotes glial infiltration, and a diffusible one required for efficient phagocytosis (Fig. 8). This model is consistent with the “cloud model” of chemokine presentation ^27^, in which GAGs maintain a local “cloud” of ligand composed of both bound and “free” molecules, thereby enabling proper receptor activation. In this framework, the inability of Orion-CD2 to support normal phagocytosis may reflect the absence of a sufficiently diffusible pool of Orion. More broadly, based on the “cloud model”, a balance between membrane-associated by GAG-binding and free GAG-surrounding Orion could be essential for proper receptor activation and downstream phagocytosis. Our model suggests that Orion acts as a signal that is both locally concentrated and partially diffusible, enabling glia to selectively target specific axons for engulfment while efficiently digesting neuronal debris.

Furthermore, we observed Orion associated with EV, most probably exosomes. These circulating cellular compartments represent an interesting system to transport and amplify protein effects ^59^. Additionally, exosomes may protect ligands from extracellular degradation and help to target it to specific regions ^60, 61, 62^. We demonstrated that Orion also associates to GAGs to gain access to the exosome biogenesis pathway where it co-localizes with several proteins involved in this pathway as Rab GTPases, LAMP and CD63 ^63^. Moreover, we showed disruption of this pathway impairs adult γ-neuron remodeling. For this, we focused on proteins involved in the late stages of the exosome pathway such us Rab11, Rab35 or Rab39 while avoiding components associated with earlier stages, particularly those linked to endocytosis such as Rab5 proteins or ESCRT complexes. This distinction was important because previous studies have shown entry into the endocytic pathway for proteins such as Patched is compulsory for pruning ^41^, and our goal was to avoid interfering with these processes. Interestingly, mosaic analysis with a repressible cell marker (MARCM) data suggested that Rab11 was not involved in pruning ^41^, contrasting with our findings using MB driven *UAS-Rab11^DN^*, which leads to pruning defects. This phenotype may be explained by a non-cell-autonomous role of exosomes in γ neuron pruning. Together these data represent to our best knowledge the first *in vivo* indication of the involvement of exosomes in γ neuron-glia communication during MB pruning.

Several studies have demonstrated that ligands exposed on the surface of exosomes can trigger either cell migration or phagocytosis. For example, the association of Wnt proteins with the exosomal surface, observed both during *Drosophila* development and in human cells, has been shown to induce cell migration ^64^. In addition, the presence of calreticulin, a very conserved multifunctional calcium binding protein, on exosome surfaces is known to promote phagocytosis by astrocytes ^65^. The precise functional role of exosomes carrying Orion in either infiltration or phagocytosis remains unclear. Notably, for infiltration, because the potential formation of a ternary complex involving exosome-GAG-bound Orion, axonal GAGs, and glial Draper is likely constrained by spatial considerations, such a complex seems improbable. In contrast, for phagocytosis, exosome-bound Orion can readily interact with glial Draper, locally activating the phagocytic machinery. In this scenario, exosome-bound Orion would act as a general amplifier of the phagocytic machinery, whereas the freely diffusing pool of Orion within the extracellular GAG-surrounding “cloud” would provide the specific cues necessary for appropriate phagocytosis. Importantly, the existence of an additional free diffusing pool of Orion is supported by our previous work using MARCM clones ^18^ which demonstrated a non-γ-neuron-autonomous function of Orion, suggesting that it can travel between axons to exert its effects. However, we cannot exclude that this transport may also be mediated by exosomes.

This study suggests that Draper activation within a multi-component signaling platform, may depend not only on ligand binding but also on the mode of ligand presentation. In addition to spatial regulation, temporal control of Orion presentation may further contribute to the sequential activation of glial responses. Glial infiltration and phagocytosis occur in a defined order during neuronal remodeling, raising the possibility that distinct pools of Orion are generated or become predominant at different developmental stages. Such a temporal switch in ligand presentation could ensure the coordinated progression from infiltration to phagocytosis.

Our results challenge the classical distinction between diffusible and membrane-tethered signals. Rather than acting simply as a secreted or membrane-associated ligand, Orion appears to function within a range of presentation states. The coexistence of membrane-associated and more diffusible pools suggests that spatial confinement and local availability are key determinants of signaling specificity. Similar mechanisms could operate in other systems, where fine spatial and biochemical control of ligand availability might be required to elicit distinct cellular responses.

Finally, although Orion conservation in vertebrates is questionable (not identifiable through standard bioinformatic analysis because either not or only weakly conserved), its receptor Drpr is homologous to MEGF10, a receptor for C1q, expressed in vertebrate astrocytes and involved in clearance of dying cells ^66^. This suggests that the regulation of astrocyte infiltration and phagocytic signalling *via* ligand distribution, for example C1q, may constitute a conserved principle. Consequently, similar mechanisms might operate in the human brain and could provide a basis of new therapeutic strategies.

## Materials and Methods

### Orion expression and purification

#### Baculovirus preparation

Orion B (UniProtKB ID Q7KVT8, hereafter named Orion) and Orion B mutant versions in its GAG1: K243A, R244A, R247A, K250A, K253A and GAG3: R548A, R549A, K551A, R553A binding motifs named Orion Mutant GAG1 and Orion mutant GAG3 respectively, were expressed in *Spodoptera frugiperda* pupal ovarian cells 21 (Sf21) using the baculovirus expression vector system. The coding sequence of the endoplasmic reticulum chaperone BiP (UniProtKB ID P29844, residues 1-18) secretion signal, in frame with Orion residues 22-646 followed by a TEV cleavage site, an 8-His tag and a Twin-Strep-tag was inserted between *BamHI* and *EcoRI* sites of the pFastBac1 plasmid by GeneCust (Boynes, France). Bacmids preparation were performed as already described ^28^. Briefly, DH10-EMBacY cells were transformed using each construct and streaked for blue-white screening on 50 µg/mL kanamycin, 10 µg/mL tetracyclin, 10 µg/mL gentamycin, 1 mM IPTG, 200 µg/mL Bluo-Gal LB-Agar plates. White colonies were restreaked to ensure proper phenotype before growing a single colony for each construct in LB supplemented with Kan/Gent. Bacmids were extracted through alkaline lysis and used to transfect Sf21 insect cells with TransIT-Insect Transfection Reagent (Mirus Bio). Cell infection was followed thanks to the expression of the Yellow Fluorescent Protein encoded in the Bacmid and the virus of first generation (V0) was harvested 72 h post-infection. This virus was amplified by infecting another batch of Sf21 cells and high-density virions (V1) were collected by centrifugation two days after proliferation arrest. V1 solutions were stored in the dark at 4°C until further use.

#### Orion and Orion-His-Strep expression and purification

The day of infection, 500 mL of Sf21 cells culture was adjusted to a density of 1×10^6^ cells/mL and infected with 1-7.5 mL of V1. Cell density was monitored every day until proliferation arrest. 48 h after that point, cell supernatant was harvested by centrifugation for 20 min at 500×*g* at 4°C and supplemented with 3 tablets of complete EDTA-free protease inhibitors (Roche) followed by filtration on a 0.45-µm filter. Two strategies were used to purify Orion with or without its C-terminus tags.

For Orion-His-Strep, the conditioned medium was loaded onto a 5-mL Strep-Tactin XT 4Flow column (IBA Lifesciences) followed by washing in 20 mM HEPES, 500 mM NaCl, pH 7. Finally, Orion-His-Strep was eluted in the same buffer supplemented with 75 mM biotin (Sigma, B4501) and underwent cycles of concentration/dilution on a 10-kDa regenerated cellulose centrifugal device (Merck, Amicon) to remove free biotin.

For Orion, the conditioned medium was loaded on a 5-mL HisTrap FF column (Cytiva). The resin was washed with 80 mL of 25 mM HEPES, 500 mM NaCl, 10 mM imidazole, pH 7.4, before elution with a linear imidazole gradient (10 to 500 mM). The C-terminus tags were removed by adding 1 mg of His-tagged TEV protease and dialyzing the sample overnight *versus* 25 mM HEPES, 150 mM NaCl, pH 7.5. A second loading step was performed on the same column to remove the His-tagged TEV protease and uncleaved Orion-His-Strep. Finally, fractions containing tag-free Orion were concentrated to 500 µL and injected on a size exclusion chromatography column (Cytiva, Superdex 200 Increase 10/3000 GL) pre-equilibrated in 25 mM HEPES, 150 mM NaCl, pH 7.5 and elution fractions were pooled and flash freezed in liquid nitrogen.

### Three-dimensional model of Orion and Heparin docking

AlphaFold3 ^29^ was used to generate tridimensional models of Orion (residues 22-646) and its mutants. Electrostatic potential was calculated with UCSF ChimeraX ^30^ version 1.10.

Potential HS binding sites in Orion was analyzed with the ClusPro 2.0 server ^31^ by docking an HS tetrasaccharide (IdoA2S-GlcNS6S)2 onto Orion after deletion of the residues 22-40, predicted to be disordered.

### Biotinylation of Heparan Sulfate

HS purified from porcine intestinal mucosa (Celsus Lab, HI-11098) was biotinylated at its reducing end as already described ^32^. Briefly, HS (5 mg/mL) was incubated in 100 mM acetate buffer (pH 4.8) containing 10 mM aniline and 12.5 mM alkoxyamine-PEG4-biotin (Thermo Fisher Scientific, 26137) at 37°C for 16–24 h. Excess biotin reagent was removed by chromatography using two HiTrap desalting columns (Cytiva) connected in series and pre-equilibrated in water.

### Surface Plasmon Resonance (SPR) Analysis

SPR experiments were performed on a BIAcore T200 instrument (Cytiva). Streptavidin was immobilized at a flow rate of 5 µL/min on a CM4 Series S sensor chip *via* amine coupling. The flow cells were activated with a mixture of 200 mM N-ethyl-N′-(diethylaminopropyl)-carbodiimide and 50 mM N-hydroxysuccinimide. Streptavidin (Sigma, S4762) was injected at 100 μg/mL in 10 mM acetate buffer, pH 4.2 until approximately 2000 response units (RU) were reached. One flow cell was used as a reference control. Biotinylated HS was then injected over the measurement flow cells to achieve immobilization levels of 40 RU for binding analysis or 130 RU for competition assays.

Binding assays were conducted in HBS-P buffer (10 mM HEPES, 150 mM NaCl, 0.05% surfactant P20, pH 7.4) at a flow rate of 30 μL/min. For kinetic analysis, recombinant untagged Orion was injected at concentrations ranging from 25 to 200 nM for 4 min, followed by a dissociation phase (8 min). The surfaces were regenerated after each cycle by injecting 2 M NaCl for 30 s. Sensorgrams were obtained by subtracting the signal from the reference flow cell. The binding curves were evaluated with the Biaevaluation 3.2.1 software and the interaction data were analyzed by fitting both association and dissociation phases.

For competition assays, Orion (75 nM) was pre-incubated, either with unlabeled HS (CelsusLab, HI-11098), chondroitin sulfate A (Sigma, C9819), D (Seikagaku, 400676), or E (Amsbio, AMS.CSR-NACS-E2.SQC-10) at 10 μg/mL for 30 min at room temperature before injection. The mixtures were then injected across the HS channel, as above described.

### Glypican-1 purification and interaction with Orion

The coding regions of soluble human glypican-1 (GPC1, UniProtKB ID P35052) or its mutant lacking all the GAG attachment sites GPC1 ΔGAG (S486A, S488A, S490A) were transiently transfected in HEK 293F cells. The cells were incubated at 37°C under agitation, supplemented with 8% CO2. Four days later, the cell supernatant was recovered from which GPC1 was purified by affinity chromatography and biotinylated. The interaction between GPC1 and Orion-His-Strep or its mutants (Orion Mutant GAG1 and Orion Mutant GAG3) was assayed by overlay dot blot. For that purpose, 5 to 200 ng of GPC1 diluted in 30 mM Tris, 250 mM NaCl, pH 7.5, were blotted onto a 0.2-µm nitrocellulose membrane (Cytiva, Amersham Protran 0.2 NC). The membrane was blocked for 1h30 in Tris buffered saline supplemented with 0.1% Tween-20 (TBS-T) and 3% BSA (Sigma, A7030), followed by incubation with 3 µg Orion-His-Strep (wild type, Orion Mutant GAG1, or Orion Mutant GAG3) diluted in the same solution for 1 h. The membrane was washed 3 times for 5 min in TBS-T, then incubated with anti-polyHis antibody coupled to horseradish peroxidase (Sigma, A7058) in TBS-T + 3%BSA for 1 h. The membrane was finally washed 3 times for 5 min in TBS-T and revealed using SuperSignal WestPico PLUS (ThermoScientific) and a ChemiDoc MP system (BioRad).

### Drosophila stocks

All crosses were performed using standard culture medium at 25 °C. Except were otherwise stated, alleles have been previously described (http://flystocks.bio.indiana.edu). *orion^null^* mutants (*orion*^1^ and *orion^ΔC^*) and *UAS-orion-CD2 myc* were produced in previous studies in our laboratory ^18, 19^; *UAS-orionB-CD2-mIFP* (named here *UAS-orion-CD2*), *UAS-orionB-GFP* and *UAS-MApHS* were a gift from Chun Han (Weill Institute, Cornell University, NY, USA), *UAS-Rab7-GFP* was provided to us from Chun Han and comes from Hugo Bellen (Baylor College of Medicine, University of Houston, TX, USA), *UAS-orion-GPI* was generated in this work. We used two GAL4 lines: *201Y-GAL4* expressed in MB γ-neurons and *Lpp-GAL4* expressed in fat body. *glypican ^ΔGAG^* was from Hiroshi Nakato (University of Minnesota, MN, USA).

### UAS Constructs

*Orion-GPI* nucleotide sequence contained the complete *Orion B*-cDNA sequence without the stop codon followed be the *Myc* sequence: ATGGAGCAAAAGCTCATTTCTGAAGAGGACTTGAATGAAATGGAGCAAAAGCTCATTTCTGAAGAGGACTTGAATGAA and the Dally GPI domain GGGTCCAACCCACTGGAAGCGACCGCGACGTGGATGCTCCTCACTCTAGTCACAATGCTCTTTAGTAGCTGTAGTTAA (corresponding to amino acids 602-626 of Dally). The resulting *orion-myc-GPI* cDNA produced at GenScript (Piscataway, NJ) in the pcDNA3.1-C-(k)DYK vector. Amplified cDNA was processed for pENTR/D-TOPO cloning (Thermo Fisher Scientific, K240020) and constructs were subsequently sequenced (Genewiz, France). We used the Gateway LR clonase enzyme mix (Thermo Fisher Scientific, 11791019) to recombine the inserts into the destination UAS vector pJFRC81-GW-2× Myc (Lee G. Fradkin, unpublished), which was generated from pJFRC81-10XUAS-IVS-Syn21-GFP-p10 (Addgene, plasmid 36462 deposited by G. Rubin ^33^ by replacing the GFP ORF with a Gateway cassette adding on a C-terminal 2X Myc tag. Transgenic fly lines (*UASs*) were generated at BestGene and the transgene was inserted into the *attP* site VK00027 in 89E11. All the crosses involving the *UAS-GAL4* system were performed at 25 °C.

### Brain dissection and immunostaining

For adult brains, *Drosophila* heads were fixated for one hour in formaldehyde (FA) 3,6% in phosphate-buffered saline (PBS) before brains were dissected in PBS 1X. Larvae and pupae brains were directly dissected in PBS 1X and then fixated for 15 min in FA 3.6%. After extraction, brains were treated for immunostaining techniques as previously described ^18^. Antibodies were used at adapted concentrations: mouse monoclonal anti-Drpr (8A1, DSHB) 1/300; mouse polyclonal anti-myc (9E10, Abcam) 1/300 and mouse monoclonal anti-CD2 (MCA154R, Bio-Rad) 1/250. For detection, Cy3 mouse conjugated to goat secondary antibodies (Jackson ImmunoResearch) was used at 1/300 concentration.

### Microscopy and image processing

Images were acquired using a Leica SP8 laser scanning confocal microscope (software: Leica Application Suite X ‘LAS X’) and a Zeiss LSM980 confocal microscope (software: Zeiss ZEN) with AiryscanII processing, each equipped with a x40 PLAN apochromatic 1.3 oil-immersion differential interference contrast objective lens. The oil type used was Immersal 518F. Microscopes were available at room temperature at the MRI platform in Institute of Human Genetics in Montpellier (France). For each experiment, images were taken using the same confocal settings.

### Hemolymph extraction

Eighty wandering larvae were PBS washed and used for one replicate. The larval hemolymph was extracted by puncture near the mouth hooks and introduced in a 0.5 mL perforated Eppendorf tube. After centrifugation at 3000 rpm, for 30 sec at 4°C, hemolymph was collected in 2.5 µL of a buffer containing: 1 mM phenylthiourea (PTU), protease inhibitor cocktail and 1 mM phenylmethylsulfonyl fluoride (PMSF), to prevent melanization. Supernatants were then centrifuged at 1000 rpm, for 10 min at 4°C, and then at 3000 g for 15 min. Two µL of this hemolymph extract was kept as a control.

### Extracellular vesicle (EV) isolation

Two different procedures were used for EV isolation: column purification and ultracentrifugation. For EV isolation by column purification, we used the SmartSEC™ Mini EV Isolation System kit (System Bioscience: SBI). The two collected EV fractions were then precipitated to collect protein pellets. Each sample was mixed with methanol (4:1 sample volume ratio) and chloroform (1:1), then water (3:1). After 5 min centrifugation at maximum speed, the aqueous phase was discarded. The tubes were then inverted several times after the addition of methanol (4.35:1). Five min centrifugation at maximum speed allowed the extraction of a protein pellet, which was dried before use. The pellets were finally resuspended with 20 µL Laemmli buffer 2X containing β-mercaptoethanol. Samples were denatured for 5 min at 95°C before being stored at -20°C.

For EV isolation by ultracentrifugation, we adapted a previous published protocol ^34^. Ten µL of haemolymph were collected as previously and immediately diluted in 500 µL of 1X PBS supplemented with PTU (0.1 µg/mL). Samples were first centrifuged at 750 g for 15 min at 4°C. The supernatant was then collected and centrifuged at 1500 g, for 10 min at 4°C. The resulting supernatant was then centrifuged at 14000 g for 35 min. At this step the supernatant (SN) and pellet (P) were collected and stored separately for further analysis. The SN fraction was then centrifuged at 100,000 g, for 2 h at 4°C and the pellet resuspended in a 20 µL mix containing 1X PBS, 5 µL of RIPA, and 4X Laemmli buffer containing β-mercaptoethanol. The P fraction was washed in PBS, centrifuged at 14000 g for 35 min, and resuspended as the SN pellet. All samples were denatured for 5 min at 70° C before being stored at -20° C.

### Western Blot

Proteins were separated on a 10% agarose gel (Mini-PROTEAN TGX precast gels, Bio-Rad) and transferred to a PVDF membrane (Merck). The membrane was blocked in 5% milk diluted in 1X PBS-Tween 0,1% (an hour minimum) before primary antibody addition overnight at 4 °C. The next day, after three 10 min washes in PBS-Tween 0.1%, the membrane was incubated in 5% milk diluted in 1X PBS-Tween 0,1% with the secondary antibody for an hour (minimum) at room temperature and then washed three times with PBS-Tween 0.1%. Detection was performed using ChemiDoc (Bio-Rad). Western blot antibodies were used at the following dilution: rabbit monoclonal anti-myc (71D10, Cell Signaling) 1/1000, mouse anti-α-Tubulin (12G10, DSHB) 1/500, peroxidase-conjugated AffiniPure Goat anti-mouse IgG 1/10000 (Jackson ImmunoResearch), and mouse monoclonal anti-GFP 1/1000 (A11120 (3E6) Invitrogen).

### 3D Modeling and quantification of glial infiltration

Entire image stacks were used for analysis with the 3D modeling software IMARIS (v10.0). The confocal microscopy images were converted into an IMS file for analysis using the IMARIS Converter software (v10.0). A first surface, based on the green fluorescence of the image (GFP), was created to model the total surface of the MB. A mask was created from this surface and applied to the red fluorescence (Cy3) to isolate the surface of MB infiltrating glial cells. Fluorescence thresholds were manually set to take into account individual variability in samples and associated fluorescence. A glial infiltration score (Surface of glia inside the MB/Surface of the MB) was then calculated to normalize size and morphology of the MBs.

### Quantification of neuronal remodeling defects

Z-stacked confocal images of adult dorsal and medial MB lobes were analyzed. For the dorsal lobe, the α lobes were delineated, and their fluorescence was removed. A fluorescence threshold including only unpruned γ-neurons and axonal debris was then set, giving a surface of dorsal neuronal remodeling defects. For the medial lobe, the fluorescence of the β lobes was also suppressed. A fluorescence measurement of the adult γ lobe was performed and kept as a background value, thus allowing to isolate the surface of the debris and unpruned axons that yield a stronger fluorescence level. The areas obtained in the dorsal and medial lobes were added to have one γ neuron remodeling defect area per MB.

For adult individuals expressing dominant negative forms of proteins involved in the exosome pathway biogenesis (Fig. 7), two classes were defined: wild type (absence of debris and unpruned axons), and mutant (presence of axonal debris and/or unpruned axons).

### Quantification of protein colocalization

The quantification of protein colocalization was performed with the Fiji software (v2.16.0) and the associated JACoP (Just Another Colocalization Plugin, v2.1.4) extension. Quantifications were performed on images from confocal microscopy with an Airyscan process. For each MB, a square was manually defined in the medial lobe, excluding zones without fluorochrome (black) throughout the entire thickness of the MB. After defining the surface, fluorescences were split to be analysed with JACoP. Fluorescence thresholds were set manually for each fluorochrome to reduce the impact of background noise variability. The software generated randomized Pearson coefficients.

### Phagocytosis quantification

To quantify phagocytosis, confocal images taken at the same confocal settings were analysed with the Fiji software. The red phagocytosis circles were manually delineated every five stacks in the medial lobe only to obtain a global phagocytosis surface. In parallel, the surface of the MB medial lobe (yellow labelling including the red phagosomes inside), was measured in the same way. To take into account the individual variability of MBs in terms of size and morphology, we calculated a surface ratio by dividing the obtained phagocytosis surface by the total surface of the γ MB medial lobe.

### Statistics

Comparison of two groups expressing a quantitative variable was analysed using the two-sided nonparametric Mann–Whitney U test. Graphs were created using GraphPad Prism software (version 10.4.2). Statistical significance was defined as ****P < 0.0001; ***P < 0.001; **P < 0.01; *P < 0.05; ns, not significant. The sample size of each group (n) is indicated in a parenthesis.

## Acknowledgments

We thank Hiroshi Nakato (University of Minnesota, MN, USA) for *glypican^ΔGAG^ Drosophila* stock used in this study. We thank the Bloomington *Drosophila* Stock Center (Indiana University), the BioCampus RAM-*Drosophila* facility and the BioCampus imaging facility MRI (Montpellier, France). We acknowledge BestGene and Genewiz for transgene service and DNA sequencing respectively. The 1D4 anti-Fasciclin II hybridoma and the 8A1 anti-Drpr monoclonal antibodies were obtained from the Developmental Studies Hybridoma Bank, created by the NICHD of the NIH and maintained at the University of Iowa, Department of Biology, Iowa City, IA 52242. We acknowledge Martin Pelosse for support in using the Eukaryotic Expression Facility at EMBL Grenoble and for the gift of DH10-EMBacY and insect cells. Part of the figures were created with BioRender.com. ChatGPT (GPT-5 mini, OpenAI) was used exclusively to improve the scientific writing and manuscript clarity.

## Fundings

C.G and C.P. were supported by grants from the Agence Nationale de la Recherche (ANR-24-CE16-5161-ExOrion and ANR-21-CE16-0033-ORIO respectively). Work was supported by the Centre National de la Recherche Scientifique (CNRS) and the Agence Nationale de la Recherche (ANR-21-CE16-0033-ORIO to J-MD. and ANR-24-CE16-5161-ExOrion to AB).

This work used the platforms of the Grenoble Instruct-ERIC center (ISBG ; UAR 3518 CNRS-CEA-UGA-EMBL) within the Grenoble Partnership for Structural Biology (PSB), supported by FRISBI (ANR-10-INBS-0005-02) and GRAL, financed within the University Grenoble Alpes graduate school (Ecoles Universitaires de Recherche) CBH-EUR-GS (ANR-17-EURE-0003).

## Notes

### Competing Interest Statement

The authors have declared no competing interest.

