## Supplemental File for "Glycosaminoglycan presentation-dependent regulation of glial responses by Orion during *Drosophila* neuronal pruning"

**Supplementary Material and methods**

**Orion protein production**

Orion sequence after secretion signal removal (bolded underlined residues represent feature typical consensus GAG binding sequences):

GNTKSASSEGWMHPSTPQMEVDALRTEYIALERALWNYLAKTANSQNNKETQIRKV
YDSHRDFDAKPQMRRTFEEHRYKILNHYEWSLLERDLIYISKLYDAYKDTFVKQSNS
VELDELAVLNLAGAILRNDNTASMPRILQEIERVMVSQTLYYRAMIVSTDQICNTMQS
AQQFIYSLYADIALTELKAYTMMEFSWMMLRVYGKGNYTQEAEIMRSEYE**KRTERTL**
**KILK**DVMLRADRIVYRCDPTKHVRGVTYDEVTRLLQGYIENEVDLNKEETCRETCDF
YQSTRSEGCFKDLYCSRQPKCSGRLYNCRFVDSDMWVCPSPQNSTRRYEFIEYEN
GRVLGQRGKCTRGTTKVDSWWRYLLWHCSYCFCLCDEEGLKSDRFFNLRDTIADV
**KRNR**VVTGLRFVKQNRIFHLQIQEGELLPRGIVNQSTLEWKPVEKYNVFDRHVKNG
VDYHKLSYEKRTIDLDDVDTDDNSFVVTGVRFRVVGTHLNLEAYYSEFDFRTGQLIR
PEYNSYWKSNDNTDVSGA**RREKLR**LSNADVSTRTIAHSIPLSRHNQYIDFTNTGLDK
DAAQSTVPFMDIQDVVSNPPVPLAGIGIYYKGRNGYGGFLGPKIITYDFMPHVQVPK
NRFENLYFQGHHHHHHHHSAWSHPQFEKGGGSGGGSGGSAWSHPQFEK

Orion Mutant GAG1 (K243A/R244A/R247A/K250A/K253A) sequence after secretion signal removal (bolded residues represent the mutated GAG1 motif):

GNTKSASSEGWMHPSTPQMEVDALRTEYIALERALWNYLAKTANSQNNKETQIRKV
YDSHRDFDAKPQMRRTFEEHRYKILNHYEWSLLERDLIYISKLYDAYKDTFVKQSNS
VELDELAVLNLAGAILRNDNTASMPRILQEIERVMVSQTLYYRAMIVSTDQICNTMQS
AQQFIYSLYADIALTELKAYTMMEFSWMMLRVYGKGNYTQEAEIMRSEYE**AATEATL**
**AILA**DVMLRADRIVYRCDPTKHVRGVTYDEVTRLLQGYIENEVDLNKEETCRETCDF
YQSTRSEGCFKDLYCSRQPKCSGRLYNCRFVDSDMWVCPSPQNSTRRYEFIEYEN
GRVLGQRGKCTRGTTKVDSWWRYLLWHCSYCFCLCDEEGLKSDRFFNLRDTIADV
KRNRVVTGLRFVKQNRIFHLQIQEGELLPRGIVNQSTLEWKPVEKYNVFDRHVKNG
VDYHKLSYEKRTIDLDDVDTDDNSFVVTGVRFRVVGTHLNLEAYYSEFDFRTGQLIR
PEYNSYWKSNDNTDVSGARREKLRLSNADVSTRTIAHSIPLSRHNQYIDFTNTGLDK
DAAQSTVPFMDIQDVVSNPPVPLAGIGIYYKGRNGYGGFLGPKIITYDFMPHVQVPK
NRFENLYFQGHHHHHHHHSAWSHPQFEKGGGSGGGSGGSAWSHPQFEK

Orion Mutant GAG3 (R548A/R549A/K551A/R553A) sequence after secretion signal removal (bolded residues represent the mutated GAG3 motif):

GNTKSASSEGWMHPSTPQMEVDALRTEYIALERALWNYLAKTANSQNNKETQIRKV
YDSHRDFDAKPQMRRTFEEHRYKILNHYEWSLLERDLIYISKLYDAYKDTFVKQSNS
VELDELAVLNLAGAILRNDNTASMPRILQEIERVMVSQTLYYRAMIVSTDQICNTMQS
AQQFIYSLYADIALTELKAYTMMEFSWMMLRVYGKGNYTQEAEIMRSEYEKRTERTL
KILKDVMLRADRIVYRCDPTKHVRGVTYDEVTRLLQGYIENEVDLNKEETCRETCDF

YQSTRSEGCFKDLYCSRQPKCSGRLYNCRFVDSDMWVCPSPQNSTRRYEFIEYEN
GRVLGQRGKCTRGTTKVDSWWRYLLWHCSYCFCLCDEEGLKSDRFFNLRDTIADV
KRNRVVTGLRFVKQNRIFHLQIQEGELLPRGIVNQSTLEWKPVEKYNVFDRHVKNG
VDYHKLSYEKRTIDLDDVDTDDNSFVVTGVRFRVVGTHLNLEAYYSEFDFRTGQLIR
PEYNSYWKSNDNTDVSGA**AAEALA**LSNADVSTRTIAHSIPLSRHNQYIDFTNTGLDK
DAAQSTVPFMDIQDVVSNPPVPLAGIGIYYKGRNGYGGFLGPKIITYDFMPHVQVPK
NRFENLYFQGHHHHHHHHSAWSHPQFEKGGGSGGGSGGSAWSHPQFEK

**Supplementary Figure Legends**

**Supplementary Figure 1- SDS-PAGE analysis of the purification of Orion.**
10% SDS-PAGE, stained with Coomassie blue, following the purification steps of Orion-His-Strep and Orion (initial purification of Orion His Trap, TEV cleavage and size exclusion chromatography Superdex 200).

**Supplementary Figure 2- Chondroitin sulfate competition on Orion binding to HS.**
Chondroitin sulfate A, D or E (CSA, CSD and CSE respectively) was preincubated with Orion for 30 min at room temperature before injection of HS surface and binding was followed by SPR (n=3). CSE inhibited the most the binding (73.8 ± 2.4 %), followed by CSA (57.2 ± 1.8 %), and CSD (27.2 ± 1.3 %). As a positive control, HS almost completely abrogated the binding (98.1 ± 2 %).

**Supplementary Figure 3- AlphaFold3 prediction of Orion structure and GAG binding sites.**
(A) Tridimensional structural model of Orion showing the α-helical (orange) and β-sheet (blue) domains. The N- (E41) and C- (F646) terminal residues are indicated. (B) Electrostatic potential mapped on the solvent accessible surface of Orion with the same orientation as in A. Values are expressed as a colour spectrum ranging from +10 kT/e (blue) to −10 kT/e (red). The dotted lines indicate the predicted GAG binding sites. (C) Close-up visualization of docking poses of an HS dp4.

**Supplementary Figure 4- SDS-PAGE analysis of the purification of Orion mutated at the GAG1 and GAG3 motifs.**

(A, B) 10% SDS-PAGE, stained with Coomassie blue, following the purification steps of Orion Mutant GAG1 and Orion Mutant GAG3. Based on the protein sequences only, the theoretical molecular weight of Orion Mutant GAG1 and Orion Mutant GAG3 are 77.9 and 78 kDa, respectively. Recovered fractions are coloured according to the construct and arrowheads indicate the protein of interest.

**Supplementary Figure 5- The association of Orion with GAGs is required for astrocytic infiltration into MB γ axon bundles at the onset of pruning.**

(A-C) Single confocal planes of 6 h APF brains expressing *201Y-GAL4*-driven *UAS-mCD8-GFP* (green). Anti-Drpr antibody (red) was used to visualize astrocytic processes. (A'-C') Anti-Drpr antibody staining (red) in which the regions corresponding to the medial lobes were delimitated with discontinuous white lines. Experiments were performed in wild type (A-A'), $orion^{\Delta C}$ (B-B'), $orion^{\Delta C}$ expressing *201Y-GAL4*-driven *UAS-orion-Mut GAG3* (C-C'). Astrocyte extension infiltration was labelled by arrows in A and C. (D) Quantification of γ lobe infiltration scores at 6 h APF by astrocytes (dorsal and medial lobe). Genotypes are listed in the supplementary Material and Methods (List of fly strains). Results are mean ± s.e.m. *n* values are indicated in a parenthesis for each condition. Replicated three times. ****$P$<0.0001 (Mann–Whitney $U$-test). Scale bar: 20μm.

**Supplementary Figure 6- Orion mutated in its GAG3 binding-domain is secreted during γ neuron remodeling.**

(A-C) Confocal Z-stacks of 6 h APF brains expressing *201Y-GAL4-driven UAS-mCD8-GFP* in combination with*: UAS-Orion-Mut GAG3* (A) and *UAS-drl* (B) in wild type; and *UAS-Orion-Mut GAG3* in $orion^1$ (C). Orion-Mut GAG3 and Drl are visualized with an anti-myc antibody (red), γ axons are visualized with GFP (green). A′-C′ (GFP) and A″-C″ (merge GFP and myc) are higher-magnification views of a single confocal plane from the corresponding square in A-C, respectively. Hole-like structures containing Orion-Mut GAG3 (red) are indicated by arrows in A', A'' and C', C''. Hole-like structures containing Drl are indicated by arrowheads in B', B''. Note the presence of high levels of secreted Orion-Mut GAG3 in these holes compared to Drl. Genotypes are listed in the supplementary Material and Methods (List of fly strains). Replicated three times. $n \geq 20$. Scale bar: 20 µm.

**Supplementary Figure 7- *dally*, *dlp*, and *orion* γ neuron mRNA expression peaks occur at similar time points during pruning**

(A-C) Data showing "*l(1)G0193*" (*orion)* (A), *dally* (B) and *dlp* (C) mRNA γ neuron-expression levels in arbitrary units (a.u.) during development. Data representations downloaded from Oren Schuldiner's laboratory's public web site (http://www.weizmann.ac.il/mcb/Schuldiner/resources). Note that the three peaks of expression occur just before the pruning process (3-6 h APF)

**Supplementary Figure 8- Orion association to plasma membrane is sufficient for astrocytic infiltration into MB γ axon bundles at the onset of pruning.**

(A, B) Single confocal planes of 6 h APF brains expressing *201Y-GAL4*-driven *UAS-mCD8-GFP* (green). Anti-Drpr antibody (red) was used to visualize astrocytic processes.

(A', B') Anti-Drpr antibody staining (red) in which the regions corresponding to the medial axon lobes were delimitated with discontinuous white lines. Experiments were performed in *orion*$^{\Delta C}$ (A-A'), *orion*$^{\Delta C}$ expressing *201Y-GAL4*-driven *UAS-orion-CD2* (B, B'). Astrocyte extension infiltration is indicated by arrows in B. (C) Quantification of γ lobe infiltration score by astrocytes at 6 h APF. Genotypes are listed in the supplementary Material and Methods (List of fly strains). Results are mean ± s.e.m. *n* values are indicated in a parenthesis for each condition. Replicated three times. ****$P$<0.0001 (Mann–Whitney *U*-test). Scale bar: 20 µm.

**Supplementary Figure 9- Orion-CD2 and Orion-GPI proteins associate to γ neuron-plasma membrane.**

(A-C) Single confocal planes of 6 h APF brains expressing *201Y-GAL4* driven *UAS-mCD8-GFP* (green) and: *UAS-orion-CD2* (A), *UAS-orion-CD2-myc* (B) and *UAS-GPI* (C). Confocal images show red Orion labelling using either anti-CD2 (A) or anti-Myc

(B, C) at the surface of $\gamma$ axons (GFP, green). Genotypes are listed in the supplementary Material and Methods (List of fly strains). $n \geq 20$ . Scale bar: 20 µm.

**Supplementary Figure 10- Plasma membrane tethered Orion induces low levels of glial phagocytosis at 2 h APF which decrease by 6 h APF**

(A-E) Single confocal planes of L3 (A-B) and 2 h APF (C-E) *orion*$^{\Delta C}$ brains expressing *201Y-GAL4*-driven *UAS-MApHS* (green/red) and *UAS-orion-myc* (A, C), *UAS-orion-CD2* (B,D) and *UAS-orion-CD2-myc* (E). Phagosomes are indicated by arrows. (F) Quantification of the phagocytosis ratio in the medial $\gamma$ lobe at different times during pupal development. Genotypes are listed in the supplementary Material and Methods (List of fly strains). Results are mean ± s.e.m. *n* values are indicated in a parenthesis for each condition. Replicated three times. *****P*<0.0001, ***P*<0.01, **P*<0.1 (Mann–Whitney *U*-test). Scale bar: 10 µm.

**Supplementary Figure 11- Orion associated to plasma membrane in combination with Orion unbound to GAGs via its GAG3 domain restores adult MB pruning**

(A, B) Single confocal planes of 6 h APF brains expressing *201Y-GAL4*-driven *UAS-mCD8-GFP* (green). Anti-Drpr antibody (red) was used to visualize astrocytic processes. (A', B') Anti-Drpr antibody staining (red) in which the regions corresponding to the medial lobes were delimitated with discontinuous white lines. Experiments were performed in *orion*$^{1}$ expressing *201Y-GAL4*-driven *UAS-orion-CD2 UAS-orion-Mut GAG3* (A-A') and *orion*$^{\Delta C}$ expressing *201Y-GAL4*-driven *UAS-orion-CD2 UAS-orion-Mut GAG3* (B-B'). Astrocyte extension infiltration was indicated by arrows. (C) Quantification of MB $\gamma$ lobe infiltration score. (D-H) Confocal Z-stacks of adult MBs expressing *201Y-GAL4*-driven *UAS-mCD8-GFP* (green) in wild type (D), *orion*$^{\Delta C}$ (E), *orion*$^{\Delta C}$; *201Y-GAL4*-driven *UAS-orion-CD2* (F), *orion*$^{\Delta C}$; *201Y-GAL4*-driven *UAS-orion-Mut GAG3* (G), *orion*$^{\Delta C}$; *201Y-GAL4*-driven *UAS-orion-CD2 UAS-orion-Mut GAG3* (H). (I) Quantification of the surface occupied by axonal debris and unpruned axons in a whole MB. Genotypes are listed in the supplementary Material and Methods (List of fly strains). Results are mean ± s.e.m. *n* values are indicated in a parenthesis for each condition. Replicated three times. ****$P$<0.0001 (Mann–Whitney $U$-test). Scale bar: 20 µm.

**Supplementary figure 12- Orion is present in exosomes and colocalizes with proteins of late-endosomes and lysosomes**

(A) Immunoblot analysis shows the presence of Orion in larval haemolymph extracts (H lanes) and exosome fractions purified by ultracentrifugation technique (P100 lanes) in *Lpp-GAL4-driven UAS-orion-myc*. $\alpha$-Tubulin was used as an internal control. Anti-Myc and anti-Tubulin antibodies were used to detect Orion and Tubulin respectively. (B-D) Single confocal planes of 6 h APF brains expressing: *201Y-GAL4-driven UAS-orion-myc* (red), *UAS-mito-GFP* (green) (B-B'), *201Y-GAL4-driven UAS-orion-myc* (red), *UAS-Rab7-GFP* (green) (C-C') and *201Y-GAL4-driven UAS-orion-myc* (red), *UAS-LAMP-GFP* (green) (D-D'). The white squares in B-D represents the region shown at higher magnification in B′-D′ in the medial $\gamma$ lobe. Colocalization of Orion (red) with exosomal proteins (green) generates yellow puncta indicated by arrows (B'-D'). Scale bars: 20 µm: B-D and 2 µm: B'-D'.

**Supplementary list of fly strains**

**Main Figures :**

**Fig.2.**

**A**. *y* $w^{67c23}$*/ Y; UAS-mCD8GFP 201Y-GAL4 /+*

**B**. *y* $w^{67c23}$ $sn^3$ $orion^1$ *FRT19A /* Y ; *UAS-mCD8GFP 201Y-GAL4 /+*

**C**. *y* $w^{67c23}$ $sn^3$ $orion^1$ *FRT19A /* Y *; UAS-mCD8GFP 201Y-GAL4 / + ; UAS-orion-B-myc / +*

**D**. *y* $w^{67c23}$ $sn^3$ $orion^1$ *FRT19A /* Y *; UAS-mCD8GFP 201Y-GAL4 / + ; UAS-orion-B-Mut GAG3-myc / +*

**Fig.3.**

**A.** *y* $w^{67c23}$*/ Y or w*/ y* $w^{67c23}$ *; 201Y-GAL4 /+ ; +; UAS-orion-B/ UAS-dally-GFP*

**B, D.** *y* $w^{67c23}$*/ Y or w*/ y* $w^{67c23}$*; 201Y-GAL4 /+ ; + ; UAS-orion-B/ UAS-dlp-GFP*

**E.** *y* $w^{67c23}$*/y* $w^{67c23}$ or *y* $w^{67c23}$*/ Y; UAS-mCD8GFP 201Y-GAL4 /+*

**F**. y $w^{67c23}$/y $w^{67c23}$ , y $w^{67c23}$/ Y or w*/Y ; UAS-mCD8GFP 201Y-GAL4 /+ ; Glypican $^{\Delta GAG}$

**Fig.4.**

**A**. y $w^{67c23}$ $sn^{3}$ $orion^{1}$ FRT19A / Y ; UAS-mCD8GFP 201Y-GAL4 /+

**B**. y $w^{67c23}$ $sn^{3}$ $orion^{1}$ FRT19A / Y ; UAS-mCD8GFP 201Y-GAL4 /+ ; UAS-orion-CD2-mIFP /+

**C**. y $w^{67c23}$ $sn^{3}$ $orion^{1}$ FRT19A / Y ; UAS-mCD8GFP 201Y-GAL4 /+ ; UAS-orion-CD2-myc /+

**D**. y $w^{67c23}$ $sn^{3}$ $orion^{1}$ FRT19A / Y ; UAS-mCD8GFP 201Y-GAL4 /+ ; UAS-orion-GPI-myc /+

**Fig.5.**

**A**. y $w^{67c23}$/ Y or y $w^{67c23}$/ w* ; 201Y-GAL4 /+ ; UAS-MApHS / +

**B**. y w* $orion^{\Delta C}$ / Y ; 201Y-GAL4 /+ ; UAS-MApHS /+

**C**. y w* $orion^{\Delta C}$ / Y ; 201Y-GAL4 /+ ; UAS-orion-B-myc / UAS-MApHS

**D**. y w* $orion^{\Delta C}$ / Y ; 201Y-GAL4 /+ ; UAS-orion-B-Mut GAG3-myc / UAS-MApHS

**E**. y w* $orion^{\Delta C}$ / Y ; 201Y-GAL4 /+ ; UAS-orion-CD2-mIFP / UAS-MApHS

**F**. y w* $orion^{\Delta C}$ / Y ; 201Y-GAL4 /+ ; UAS-orion-CD2-myc / UAS-MApHS

**Fig.6.**

**A**. y w* $orion^{\Delta C}$ / Y ; 201Y-GAL4 /+ ; UAS-orion-CD2-mIFP / UAS-MApHS

**B**. y w* $orion^{\Delta C}$ / Y ; 201Y-GAL4 /+ ; UAS-orion-CD2-mIFP UAS-orion-B-Mut GAG3-myc / UAS-MApHS

**D.** y $w^{67c23}$/ Y; UAS-mCD8GFP 201Y-GAL4 /+

**E.** y $w^{67c23}$ $sn^{3}$ $orion^{1}$ FRT19A / Y ; UAS-mCD8GFP 201Y-GAL4 /+

**F.** y $w^{67c23}$ $sn^{3}$ $orion^{1}$ FRT19A / Y ; UAS-mCD8GFP 201Y-GAL4 /+ ; UAS-orion-CD2-mIFP /+

**G.** y $w^{67c23}$ $sn^{3}$ $orion^{1}$ FRT19A / Y ; UAS-mCD8GFP 201Y-GAL4 /+ ; UAS-orion-B-Mut GAG3-myc /+

**H.** y $w^{67c23}$ $sn^{3}$ $orion^{1}$ FRT19A / Y ; UAS-mCD8GFP 201Y-GAL4 /+ ; UAS-orion-CD2-mIFP UAS-orion-B-Mut GAG3-myc /+

**Fig.7.**

**C.** *y $w^{67c23}$/ Y or y $w^{67c23}$/ w* ; UAS-mCD8GFP 201Y-GAL4 /+ ; UAS-CD63-mCherry / +*

**D.** *y $w^{67c23}$/ Y or y $w^{67c23}$/ w* ; 201Y-GAL4 /+ ; UAS-CD63-mCherry / UAS-orion-B-myc*

**E.** *y $w^{67c23}$/ Y or y $w^{67c23}$/ w* ; 201Y-GAL4 /+ ; UAS-Rab11-GFP / UAS-orion-B-myc*

**G.** *y $w^{67c23}$/ Y or y $w^{67c23}$/ w* ; UAS-mCD8GFP 201Y-GAL4 /+*

**H.** *y w67c23 $sn^3$ $orion^1$ FRT19A /* Y ; *UAS-mCD8GFP 201Y-GAL4 /+*

**I.** *y $w^{67c23}$/ Y or y $w^{67c23}$/ y w* ; UAS-mCD8GFP 201Y-GAL4 /+ ; UAS-$Rab11^{DN}$ /+*

**J.** *y $w^{67c23}$/ Y or y $w^{67c23}$/ y w* ; UAS-mCD8GFP 201Y-GAL4 /+ ; UAS-$Rab35^{DN}$ /+*

**K.** *y $w^{67c23}$/ Y or y $w^{67c23}$/ y w* ; UAS-mCD8GFP 201Y-GAL4 /+ ; UAS-$Rab39^{DN}$ /+*

**Supplementary Figures :**

**Supplementary Figure 5**

**A.** *y $w^{67c23}$/ Y or y $w^{67c23}$/ y $w^{67c23}$ ; UAS-mCD8GFP 201Y-GAL4 /+*

**B.** *y w* $orion^{\Delta C}$ /* Y ; *UAS-mCD8GFP 201Y-GAL4 /+*

**C.** *y w* $orion^{\Delta C}$ /* Y ; *UAS-mCD8GFP 201Y-GAL4 /+ ; UAS-orion-B-Mut GAG3-myc /+*

**Supplementary Figure 6**

**A.** *y $w^{67c23}$/ Y or y $w^{67c23}$/ y $w^{67c23}$ ; UAS-mCD8GFP 201Y-GAL4 /+ ; UAS-orion-B-Mut GAG3-myc /+*

**B.** *y w* $orion^{\Delta C}$ /* Y ; *UAS-mCD8GFP 201Y-GAL4 /+ ; UAS-drl-myc /+*

**C.** *y $w^{67c23}$ $sn^3$ $orion^1$ FRT19A /* Y ; *UAS-mCD8GFP 201Y-GAL4 /+ ; UAS-orion-B-Mut GAG3-myc /+*

**Supplementary Figure 8**

**A.** *y w* $orion^{\Delta C}$ /* Y ; *UAS-mCD8GFP 201Y-GAL4 /+*

**B.** *y w* $orion^{\Delta C}$ /* Y ; *UAS-mCD8GFP 201Y-GAL4 /+ ; UAS-orion-CD2-mIFP /+*

**Supplementary Figure 9**

**A.** *y $w^{67c23}$/ Y or y $w^{67c23}$/ y $w^{67c23}$; UAS-mCD8GFP 201Y-GAL4 /+ ; UAS-orion-CD2-mIFP /+*

**B.** *y $w^{67c23}$/ Y or y $w^{67c23}$/ y $w^{67c23}$; UAS-mCD8GFP 201Y-GAL4 /+ ; UAS-orion-CD2-myc/+*

**C.** *y $w^{67c23}$/ Y or y $w^{67c23}$/ y $w^{67c23}$; UAS-mCD8GFP 201Y-GAL4 /+ ; UAS-orion-GPI-myc /+*

**Supplementary Figure 10**

**A.** *y w* $orion^{\Delta C}$* / Y *; 201Y-GAL4 /+ ; UAS-orion-B-myc / UAS-MApHS*

**B.** *y w* $orion^{\Delta C}$* / Y *; 201Y-GAL4 /+ ; UAS-orion-CD2-mIFP / UAS-MApHS*

**C.** *y w* $orion^{\Delta C}$* / Y *; 201Y-GAL4 /+ ; UAS-orion-B-myc / UAS-MApHS*

**D.** *y w* $orion^{\Delta C}$* / Y *; 201Y-GAL4 /+ ; UAS-orion-CD2-mIFP / UAS-MApHS*

**E.** *y w* $orion^{\Delta C}$* / Y *; 201Y-GAL4 /+ ; UAS-orion-CD2-myc / UAS-MApHS*

**Supplementary Figure 11**

**A.** *y $w^{67c23}$ $sn^3$ $orion^1$ FRT19A* / Y ; *UAS-mCD8GFP 201Y-GAL4 /+ ; UAS-orion-CD2-mIFP UAS-orion-B-Mut GAG3-myc /+*

**B.** *y w* $orion^{\Delta C}$* / Y *; UAS-mCD8GFP 201Y-GAL4 /+ ; UAS-orion-CD2-mIFP UAS-orion-B-Mut GAG3-myc /+*

**D.** *y $w^{67c23}$/ Y; UAS-mCD8GFP 201Y-GAL4 /+*

**E.** *y w* $orion^{\Delta C}$* / Y ; *UAS-mCD8GFP 201Y-GAL4 /+*

**F.** *y w* $orion^{\Delta C}$* / Y ; *UAS-mCD8GFP 201Y-GAL4 /+ ; UAS-orion-CD2-mIFP /+*

**G.** *y w* $orion^{\Delta C}$* / Y ; *UAS-mCD8GFP 201Y-GAL4 /+ ; UAS-orion-B-Mut GAG3-myc /+*

**H.** *y w* $orion^{\Delta C}$* / Y ; *UAS-mCD8GFP 201Y-GAL4 /+ ; UAS-orion-CD2-mIFP UAS-orion-B-Mut GAG3-myc /+*

**Supplementary Figure 12**

**B.** *y $w^{67c23}$/ Y or y $w^{67c23}$/ w*; 201Y-GAL4 / UAS-mito-GFP ; UAS-orion-B-myc /+*

**C.** *y $w^{67c23}$/ Y or y $w^{67c23}$/ w*; 201Y-GAL4 / UAS-Rab7-GFP ; UAS-orion-B-myc /+*

**D.** *y $w^{67c23}$/ Y or y $w^{67c23}$/ w*; 201Y-GAL4 / UAS-LAMP-GFP ; UAS-orion-B-myc /+*

### Supplementary Figure 1

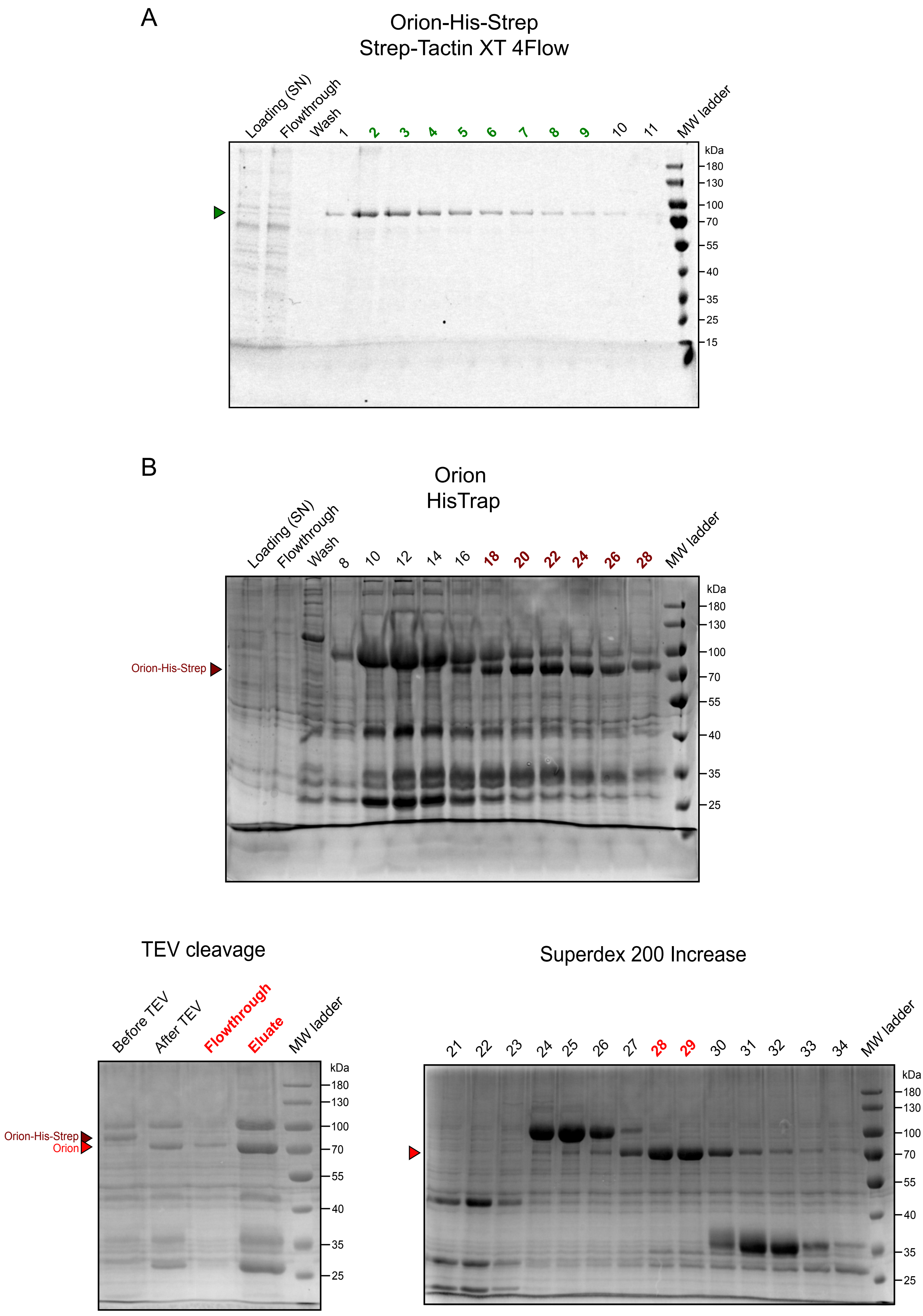

#### Supplementary Figure 2

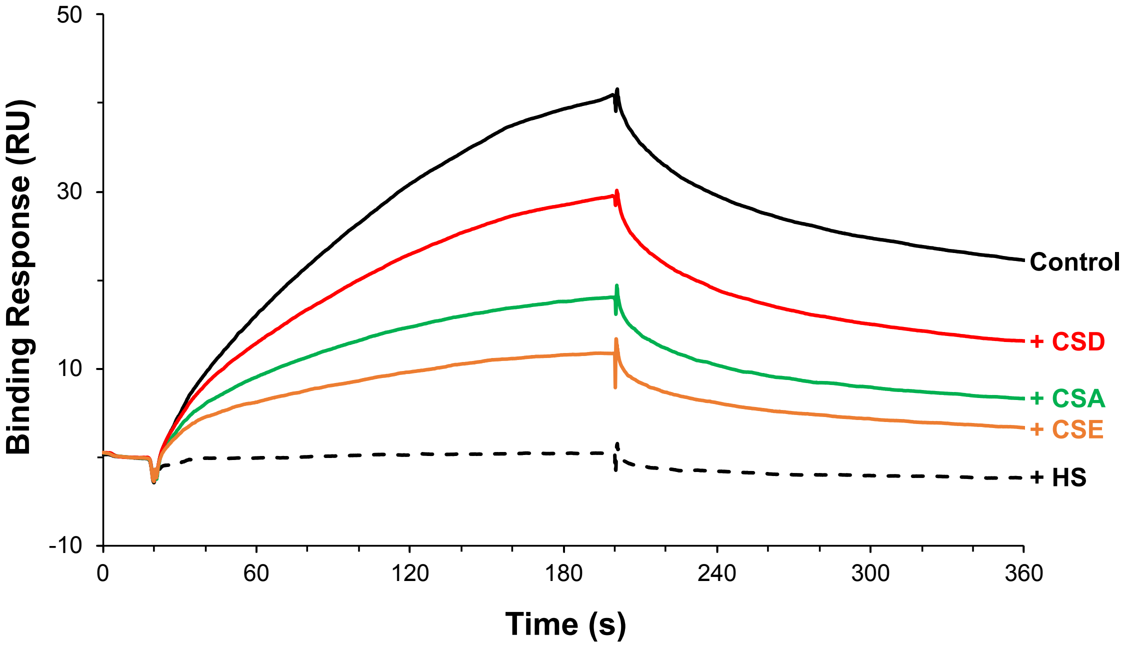

### Supplementary Figure 3

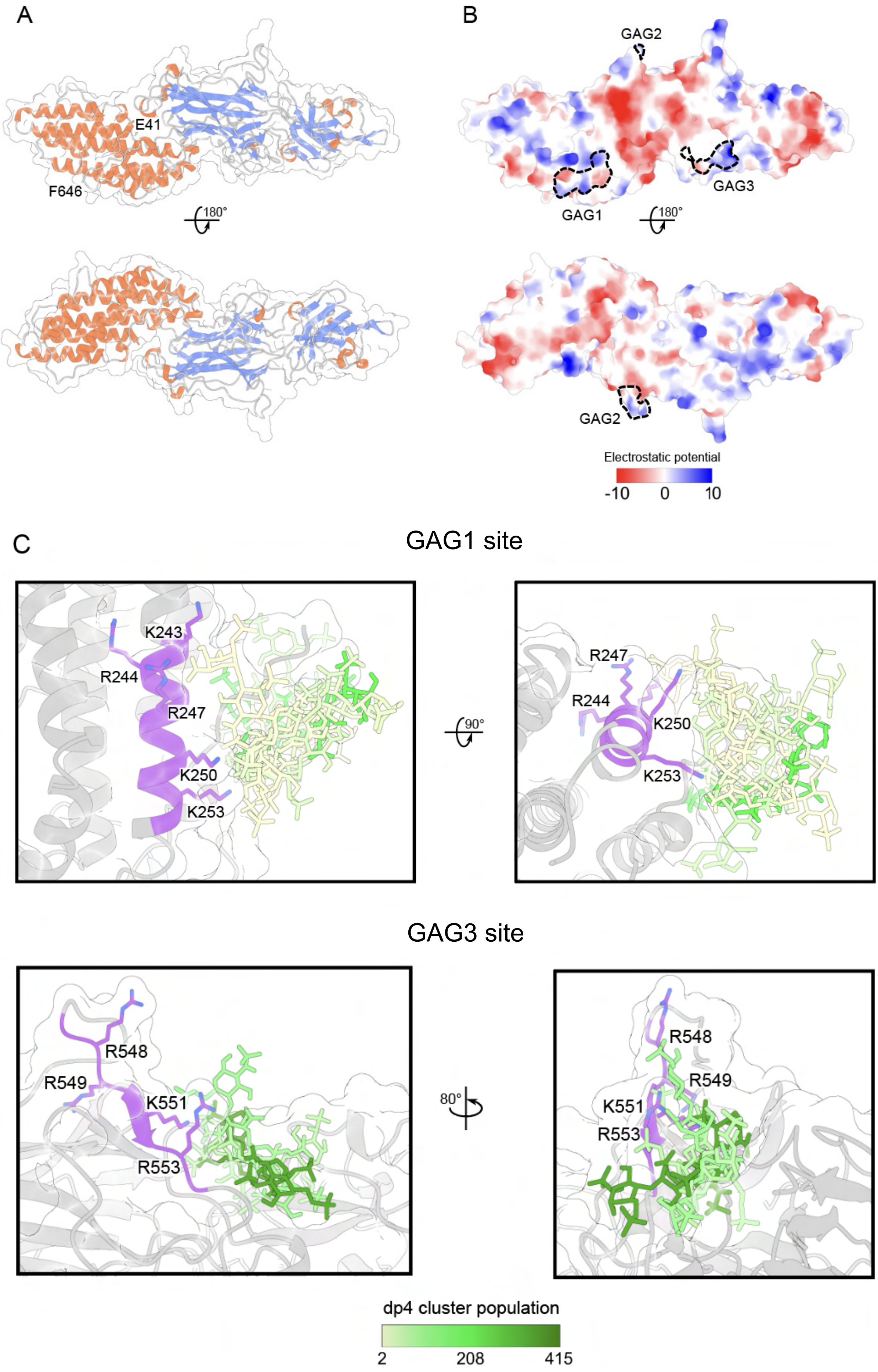

### Supplementary Figure 4

A

Orion Mutant GAG1
Strep-Tactin XT 4Flow

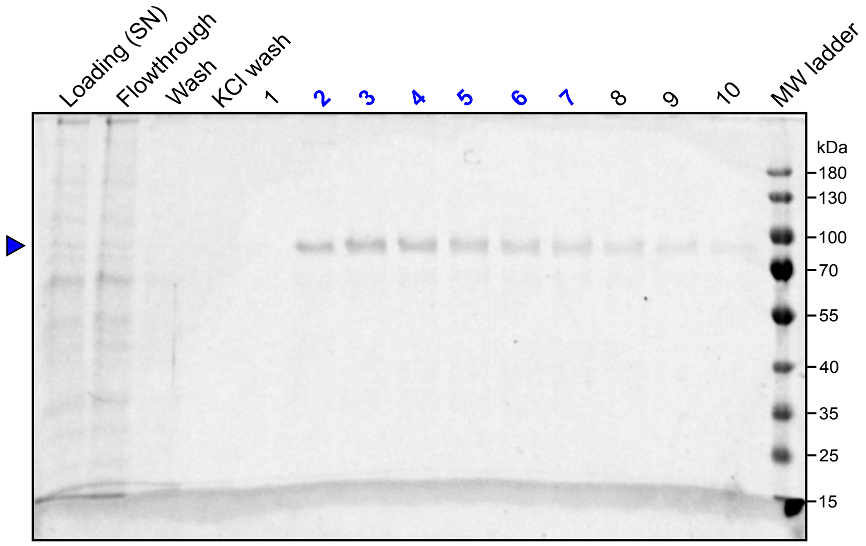


B

Orion Mutant GAG3
Strep-Tactin XT 4Flow

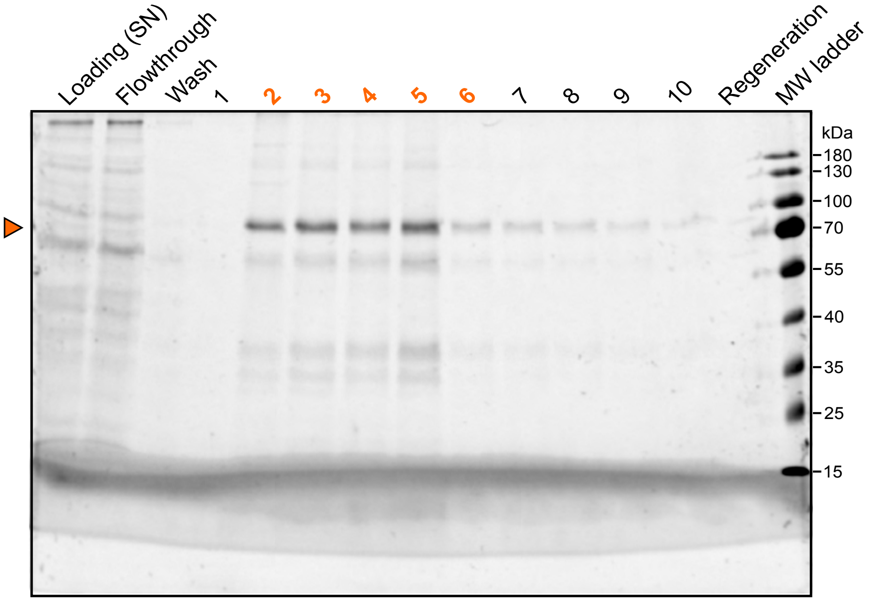

### Supplementary Figure 5

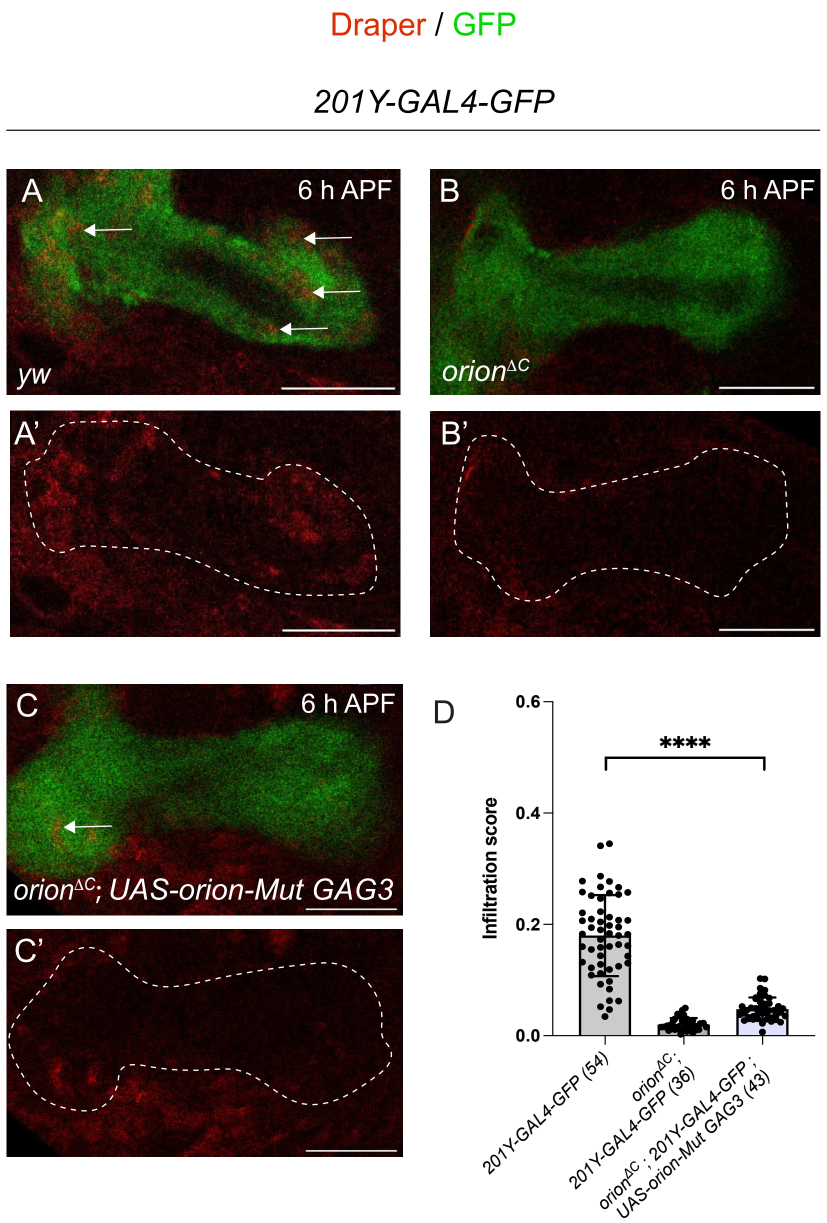

### Supplementary Figure 6

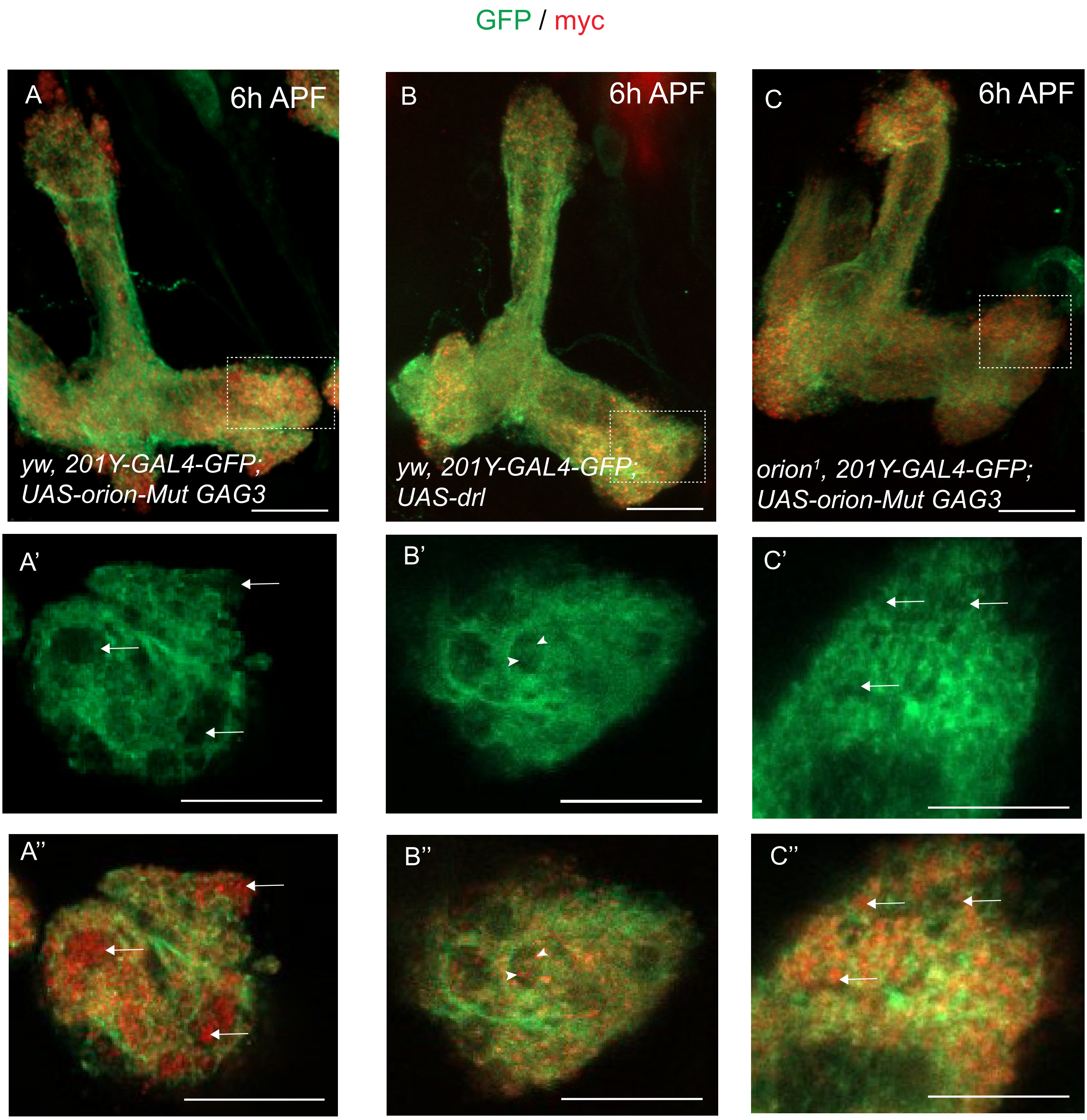

#### Supplementary Figure 7

A

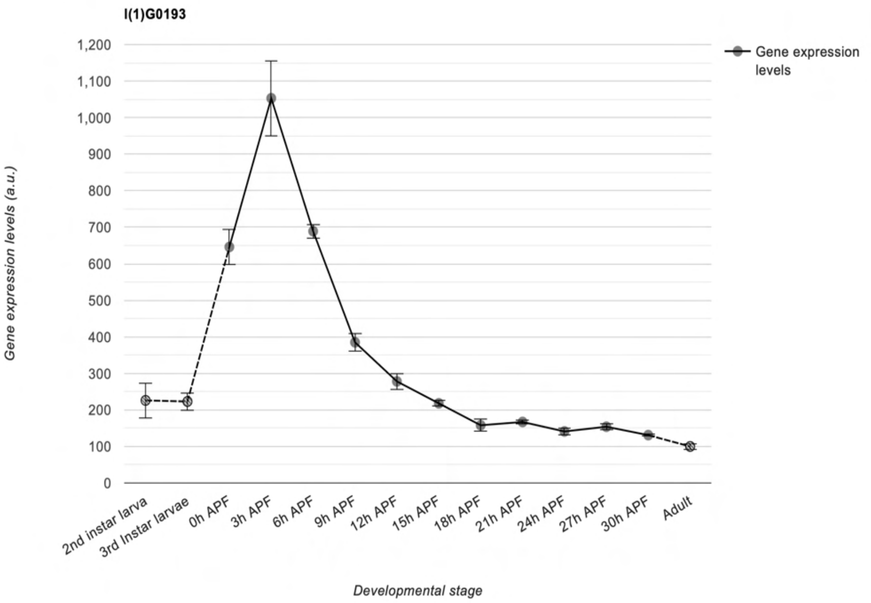


B

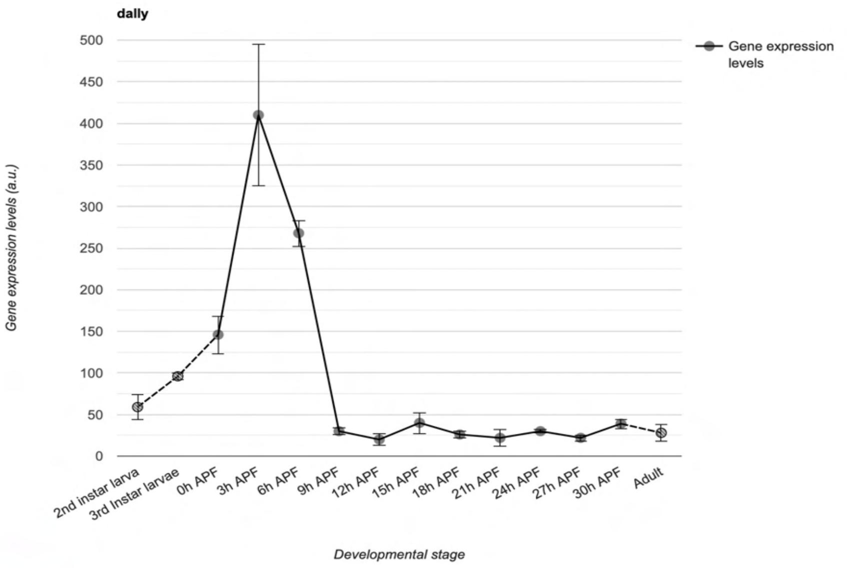


C

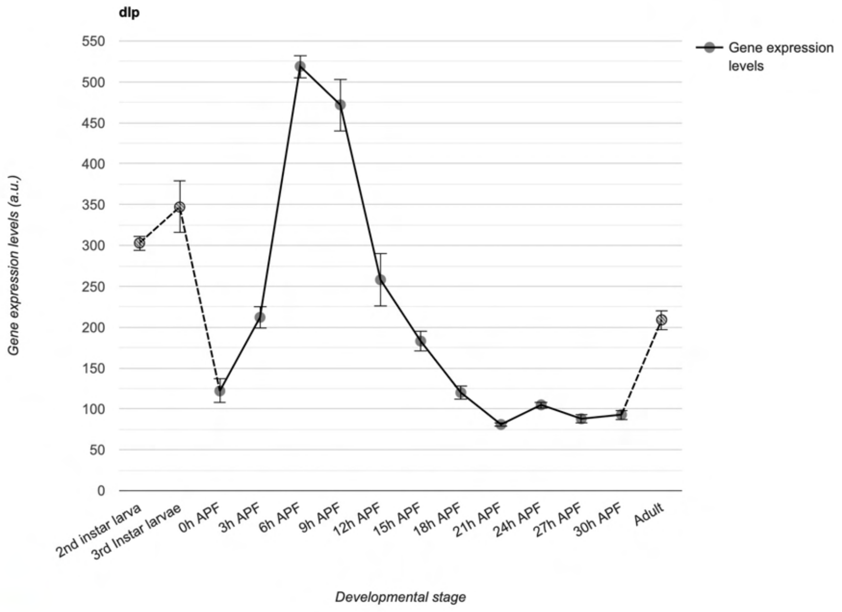

#### Supplementary Figure 8

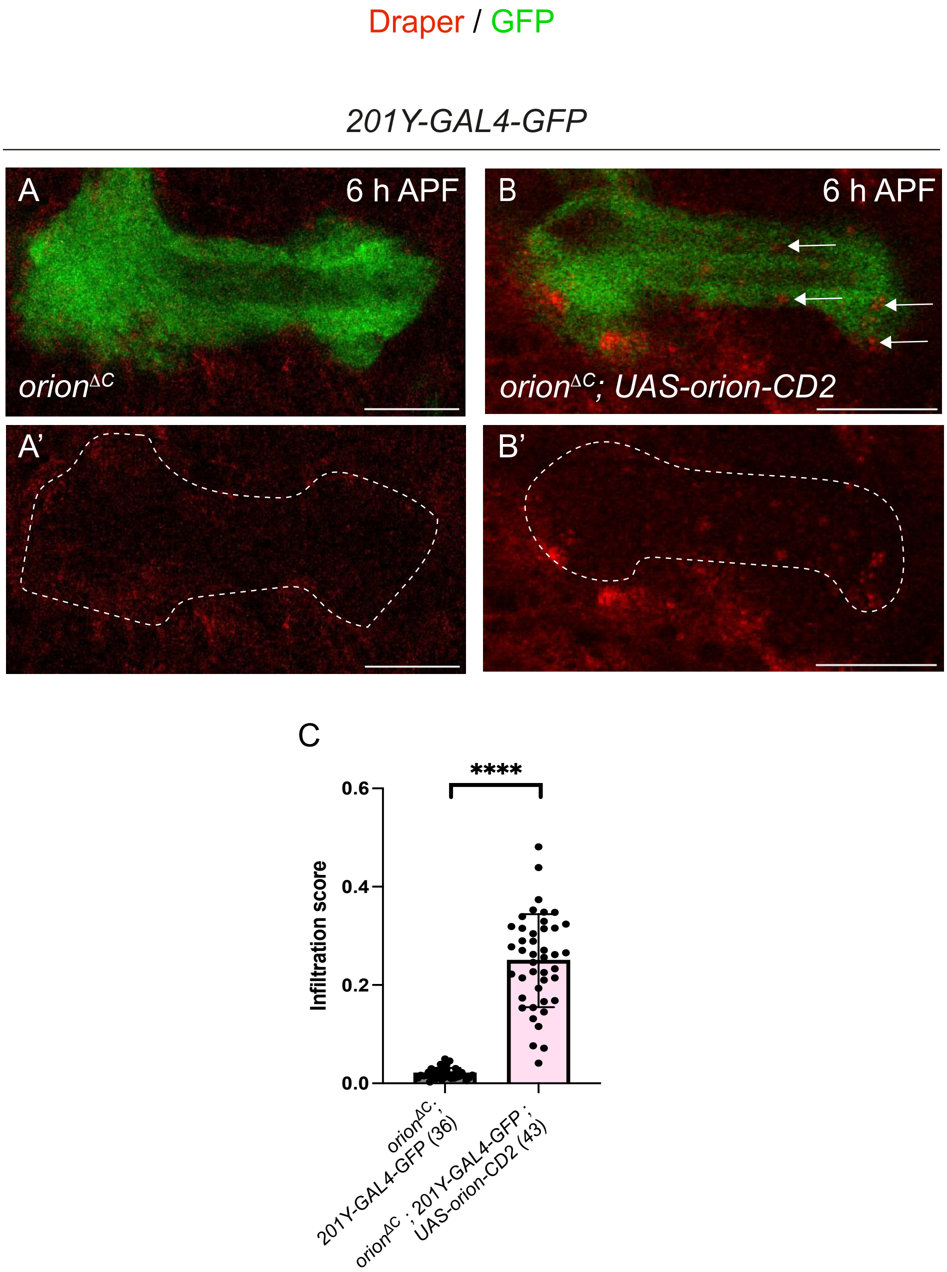

**Supplementary Figure 9**

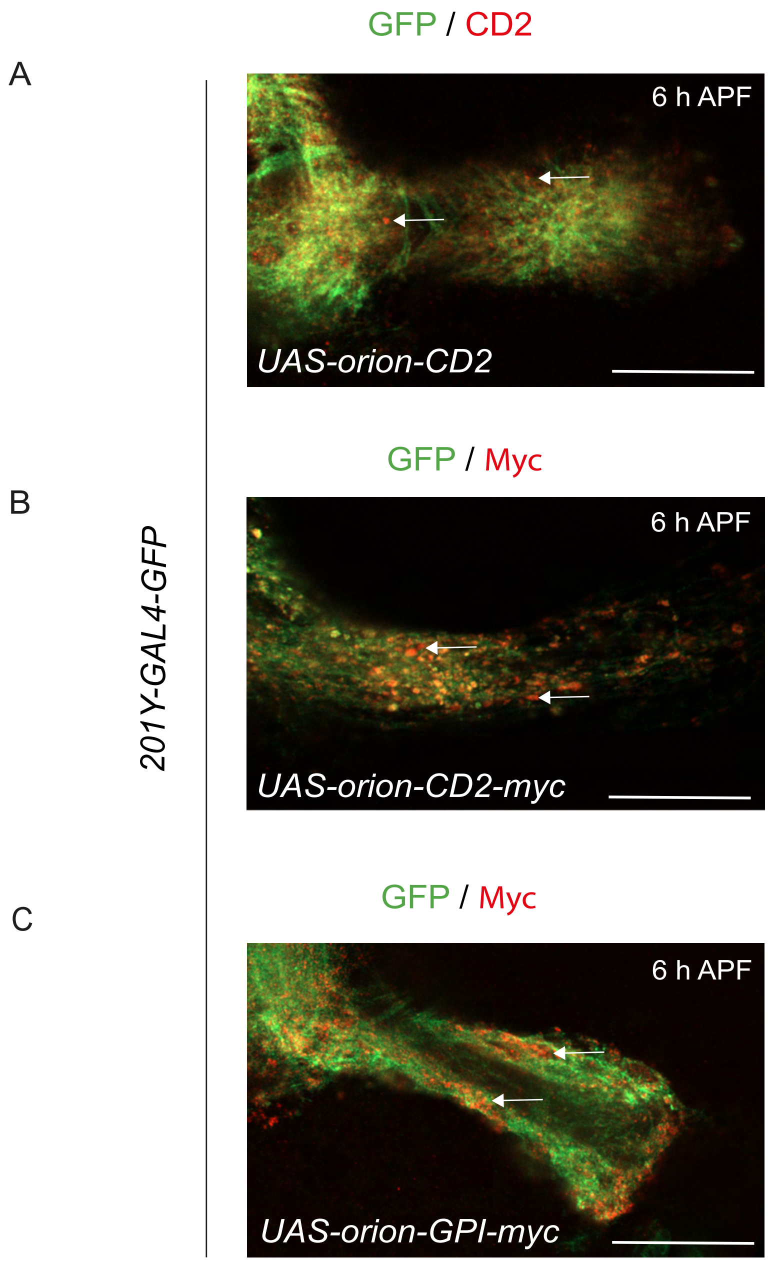

#### Supplementary Figure 10

*orion$^{\Delta C}$ ; 201Y-GAL4 ; UAS-MApHS*

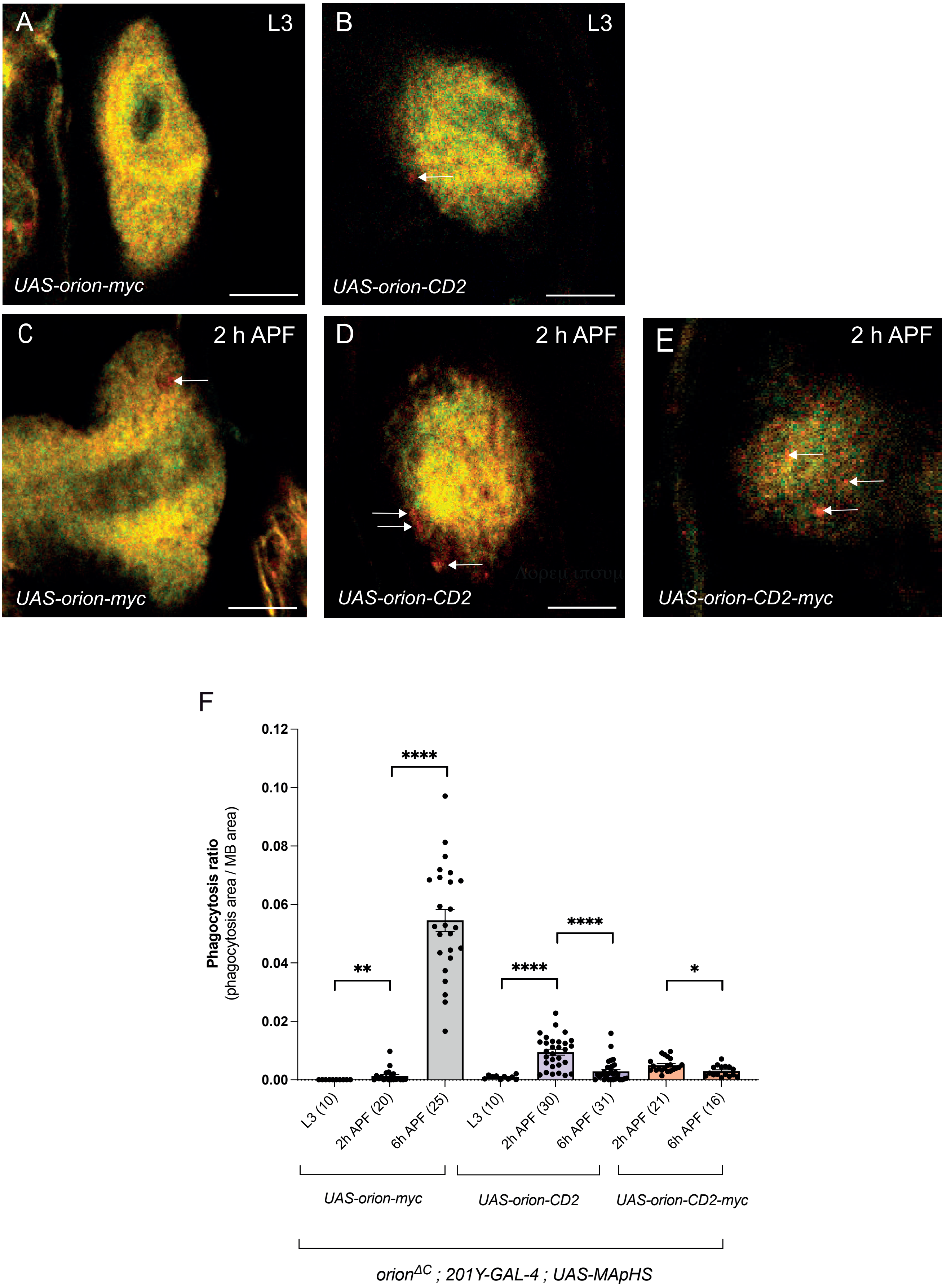

### Supplementary Figure 11

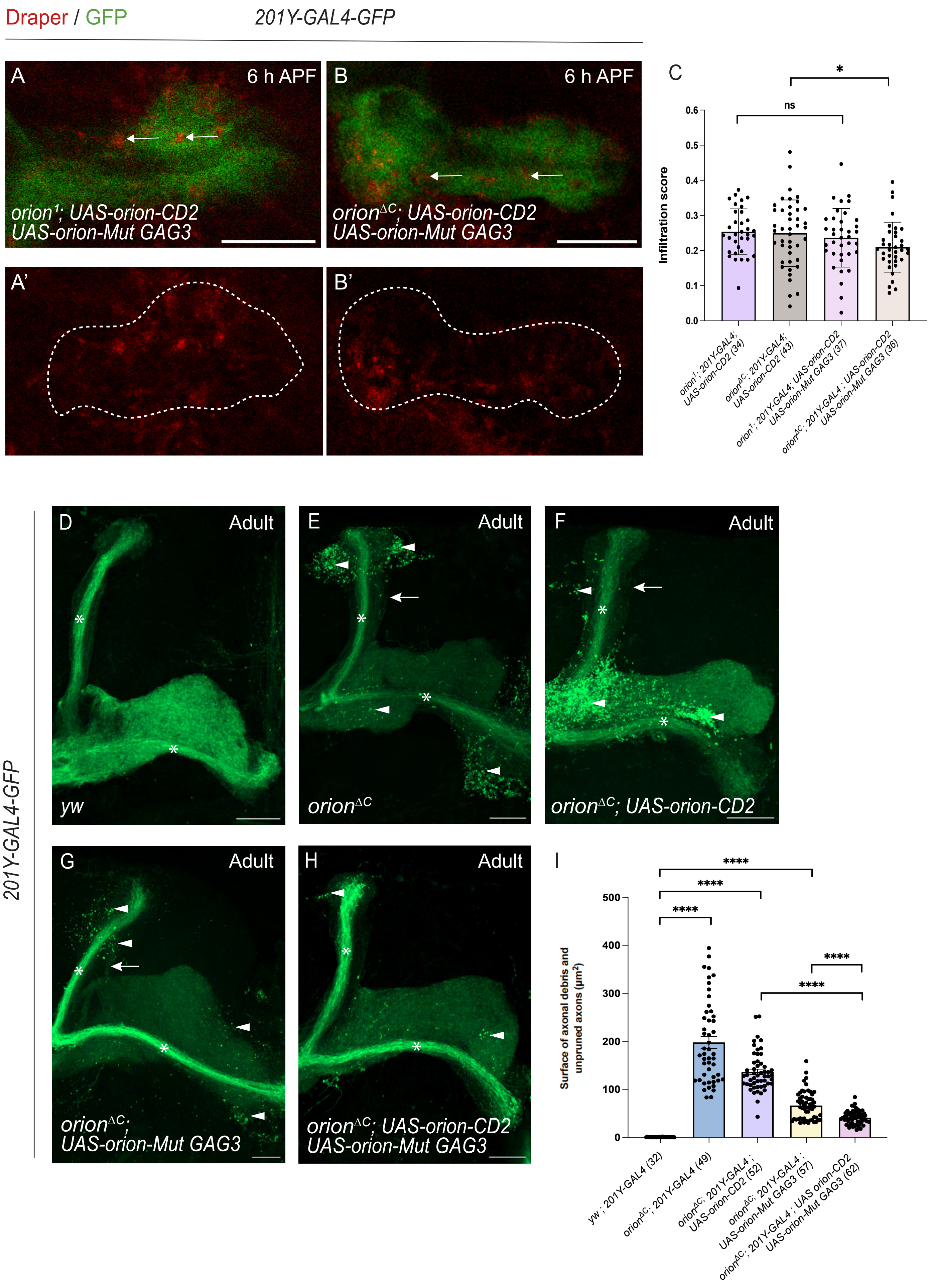

#### Supplementary Figure 12

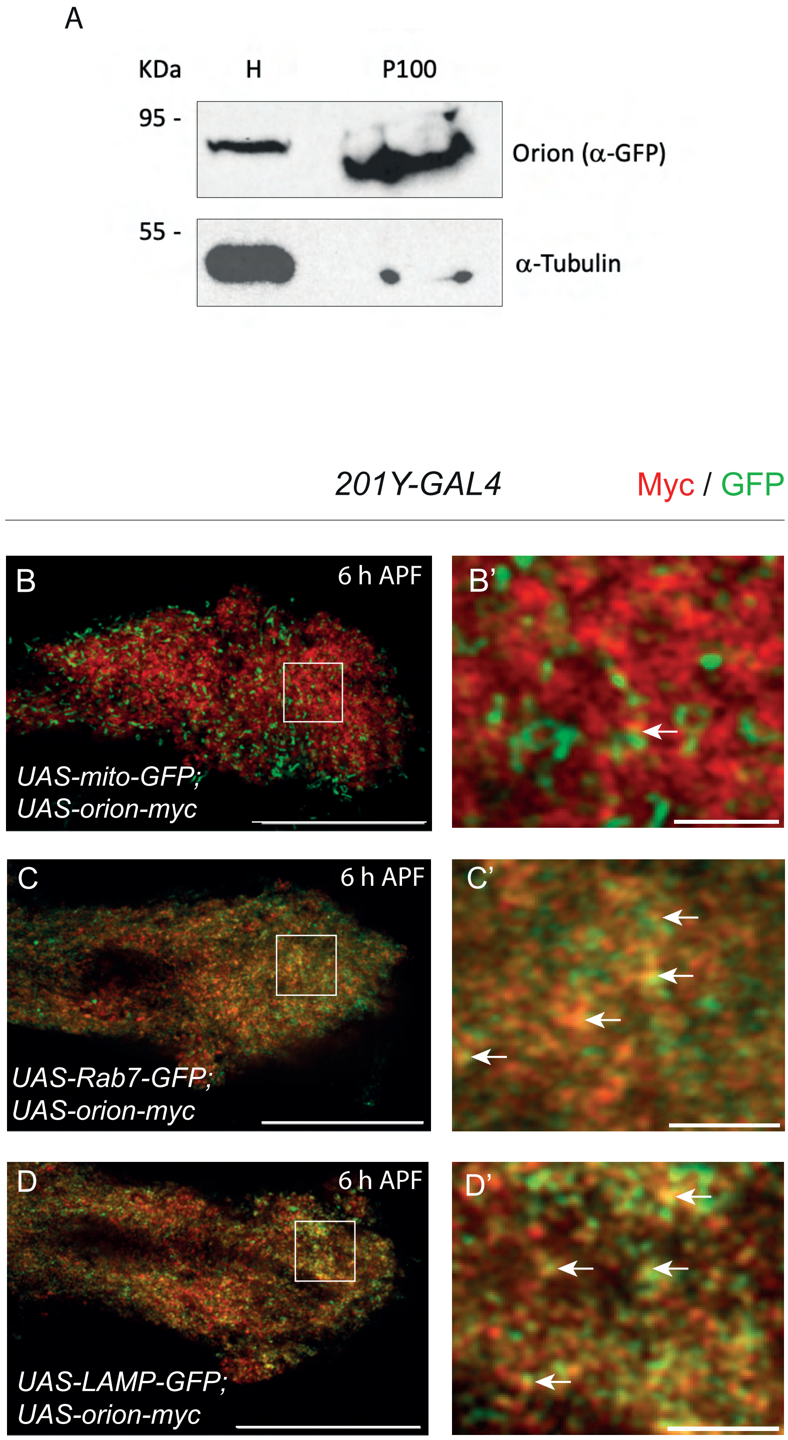
